# When is an auxotroph not an auxotroph: how budding yeast lacking *MET17* collectively overcome their metabolic defect

**DOI:** 10.1101/2023.05.18.541364

**Authors:** Sonal, Alex E. Yuan, Xueqin Yang, Wenying Shou

## Abstract

Assimilation of sulfur is vital to all organisms. In *S. cerevisiae*, inorganic sulfate is first reduced to sulfide, which is then reacted to an organic carbon backbone by the Met17 enzyme. The resulting homocysteine can then be converted to all other essential organosulfurs such as methionine, cysteine, and glutathione. This pathway has been known for nearly half a century, and *met17* mutants have long been classified as organosulfur auxotrophs which are unable to grow on sulfate as the sole sulfur source. Surprisingly, we found that *met17*Δ could grow on sulfate, albeit only at sufficiently high cell densities. We show that the accumulation of hydrogen sulfide gas underpins this density-dependent growth of *met17*Δ on sulfate, and that the locus *YLL058W* (*HSU1*) enables *met17*Δ cells to assimilate hydrogen sulfide. Hsu1 protein is induced during sulfur starvation and under exposure to high sulfide in wildtype cells, suggesting multiple functions of this gene. In a mathematical model, the low efficiency of sulfide assimilation in *met17*Δ can explain the observed density-dependent growth of *met17*Δ on sulfate. Thus, having uncovered and explained the paradoxical growth of a commonly used “auxotroph”, our findings may impact the design of future studies in yeast genetics, metabolism, and volatile-mediated microbial interactions.

## INTRODUCTION

Sulfur metabolism is vital to all organisms and produces a range of essential metabolites. Perhaps the best-known organosulfurs—sulfur-containing organic compounds—are the essential amino acids methionine and cysteine. The relevance of sulfur metabolites however extends far beyond protein synthesis. AdoMet, an activated form of methionine, serves as a universal methyl donor for nucleotides, proteins and small metabolites, and perturbations of AdoMet metabolism are implicated in liver pathologies [1]. Glutathione, generated from cysteine, is crucial for cellular redox homeostasis [2]; decreased glutathione levels leads to oxidative stress which has been implicated in ageing and neurodegeneration [3,4]. Hydrogen sulfide (H_2_S), an inorganic gas which can be produced during organosulfur metabolism, has therapeutic potential in gastrointestinal, cardiovascular, inflammatory, and nervous systems [5,6]. A deeper understanding of sulfur metabolism can thus have a wide societal impact.

Besides these critical roles in cell physiology, secreted sulfur metabolites can also influence population-level behavior in microbes. For example, in the budding yeast *Saccharomyces cerevisiae*, populations can expand the range of temperatures they can tolerate by leveraging the antioxidant properties of secreted glutathione [7]. As another example, H_2_S is thought to promote population synchrony during ultradian respiratory oscillations in aerobic continuous cultures by inhibiting the respiratory chain, and exogenous sulfide can shift the phase of these oscillations [8,9]. In fact, microorganisms release a constellation of volatile sulfur compounds [10], which have been of interest to the food industry (e.g. in optimizing wine aroma [11]), but the significance of many of these compounds to the microbes themselves is poorly understood.

Unlike humans, yeast can synthesize essential organosulfurs *de novo* by assimilating inorganic sulfates (SO_4_^2-^) from their environment. Various genes of sulfur metabolism were discovered in the budding yeast through genetic screens for organosulfur auxotrophs—mutants which can only grow when an organosulfur is supplemented and are otherwise unable to grow on sulfate [12]. These studies, together with biochemical analyses, led to the current model of sulfur metabolism in yeast [12]. In a simplified version of this model (Fig 1A), sulfate is reduced to hydrogen sulfide (H_2_S) through a series of enzymatic reactions. Hydrogen sulfide is then reacted to *O*-acetyl homoserine (OAH; Fig 1A) to form homocysteine, an organosulfur which can be interconverted to other organosulfurs (Fig 1A, purple). Some organosulfurs (e.g. cysteine) can be broken down to release H_2_S and sulfite through dedicated pathways.

**Figure 1.**
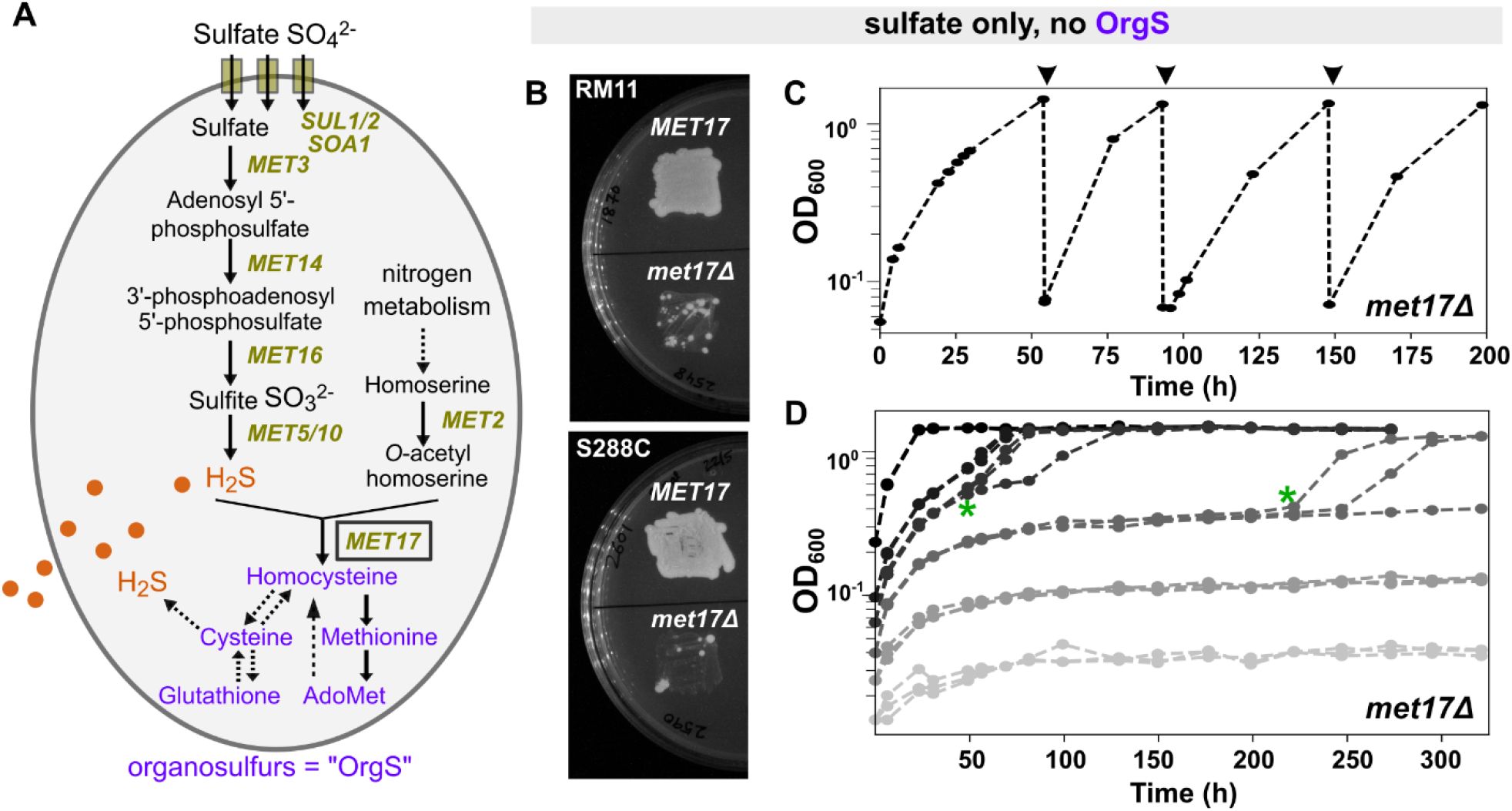
*met17*Δ cells show density-dependent growth on sulfate. **A)** Schematic summarizing sulfate assimilation in *S. cerevisiae*. Sulfate SO_4_^2-^ is reduced to sulfide S^2-^ (orange) through a series of enzymatic reactions, and Met17 then combines sulfide with a nitrogenous compound (*O*-acetyl homoserine) to generate homocysteine. All organosulfurs (purple) are interconvertible via the transsulfuration pathway (cysteine and glutathione) and the methyl cycle (methionine, AdoMet). Solid and dotted lines represent one-step and multi-step reactions, respectively. Note that according to this conventional understanding, *met17* mutants should not be able to generate organosulfurs from sulfate and therefore should require supplementation of organosulfur in their growth media. **B)** Auxotrophy of *met17*Δ is leaky. On agar plates containing synthetic minimal medium (SD: with sulfate, but without organosulfurs), patching prototrophic yeast leads to a dense lawn, while patches of *met17*Δ show papillae. Plate was imaged 5 days after patching. Prototrophic and *met17*Δ strains were WY1870 and WY2548, respectively for the RM11 background, and WY2601 and WY2590, respectively for the S288C background. **C)** *met17*Δ yeast can be repeatedly passaged on sulfate. A liquid culture of *met17*Δ in SD medium could persistently grow to saturation upon repeated dilutions (arrowheads). **D)** *met17*Δ show density-dependent growth on sulfate. Lighter and darker shades of gray represent lower and higher initial cell densities, respectively. At each starting density, three identical liquid cultures were set up in glass tubes. At higher initial cell density, all three cultures grew deterministically, while stochasticity in lag time was observed at intermediate cell densities (green asterisks). The lowest cell densities did not grow. In **B** and **C**, population growth was recorded as optical density at 600 nm (OD_600_). Each trendline represents a 7-ml culture in an 18-mm diameter glass tube with a loosely fitted plastic cap. *met17*Δ (WY2035, RM11) were grown to exponential stage in methionine-supplemented medium, and then washed and transferred to medium without methionine. WY2035 require lysine supplementation in SD medium, which was maintained throughout.

The *MET17* gene, also known as *MET15* or *MET25* [13–15], catalyzes homocysteine synthesis by reacting H_2_S with *O*-acetyl homoserine (i.e. displaying OAH sulfhydrylase activity) [16–18]. Although purified Met17 protein could also catalyze cysteine synthesis by reacting H_2_S with *O*-acetyl serine (i.e. displaying OAS sulfhydrylase activity), this activity was low [18] and unlikely to be relevant *in vivo* in *S. cerevisae* [19]. Thus, Met17 has been assigned as the sole enzyme for synthesizing homocysteine—the precursor to all other forms of organosulfurs (Fig 1A). *met17* mutants are auxotrophic for organosulfurs, and the *met17* deletion mutation (*met17*Δ) [20] is commonly used in genetic studies of yeast, for example as a background mutation in the yeast deletion library [21,22].

Surprisingly, we found that *met17*Δ yeast can in fact grow on sulfate without any organosulfur supplements, albeit in a density-dependent manner. Here, we show that density-dependence is mediated via the volatile metabolite H_2_S, which can be assimilated into organosulfurs through the *YLL058W* (*HSU1*) gene in the absence of *MET17*. A mathematical model based on low activity of Hsu1 can explain the density-dependent growth of *met17*Δ on sulfate. In Discussions, we reconcile conflicts in earlier studies, highlight considerations for studying gas-mediated microbial interactions, and speculate on the functions of *HSU1* in wildtype yeast.

## RESULTS

### *met17*Δ show density-dependent growth on sulfate

*met17* mutants were first identified in genetic screens for organosulfur auxotrophs [23] and have long been thought to be unable to grow on inorganic sulfate [12]. On synthetic minimal medium (SD) containing sulfate as the sole sulfur source, the wildtype *MET17* prototrophic strain grew a dense lawn as expected (Fig 1B). Surprisingly, in the patches of the *met17* deletion (*met17*Δ) mutant, individual colonies would appear after 1-3 days of incubation (Fig 1B), even though such patches usually remain clear for true auxotrophs. Furthermore, *met17*Δ yeast could sometimes grow to high cell densities in liquid SD medium over repeated passaging (Fig 1C). This indicated that we were not merely observing residual growth fueled by organosulfurs that the cells had accumulated during exponential growth in methionine-supplemented medium. Similar observations have since been reported by other labs [24,25], and chromatographic analyses has confirmed that organosulfur contaminants occur in negligible quantities in the commercially available yeast nitrogenous base that was used to prepare SD medium [25].

The seemingly erratic growth behavior of *met17*Δ on sulfate could potentially be explained by growth being dependent on the initial cell density in liquid cultures (Fig 1D). At high initial cell densities, growth of *met17*Δ on sulfate was deterministic. At very low initial cell densities, the cultures did not grow. Strikingly, at intermediate cell densities, growth in the cultures was stochastic: three replicate cultures started from the same parent culture would deviate in the lag time before population growth took off (Fig 1D, green asterisks). Note that the *met17*Δ cells were growing exponentially in minimal medium supplemented with methionine before being washed and transferred to minimal medium without organosulfurs. Truly prototrophic cells would exhibit similar growth behaviors independent of the initial culture density upon such a transition. Thus, *met17*Δ were not actual prototrophs but could overcome their auxotrophy at sufficiently high cell densities.

While our initial observations (Figs 1C and 1D) were made in yeast of the background RM11, we also observed a similar phenomenon in the S288C background (S1 Fig). The phenotype was however weaker: S288C *met17*Δ grew to high turbidity in minimal media but growth did not persist after dilution and required considerably higher initial cell densities in S288C as compared to RM11 (S1 Fig).

### Density-dependent growth of *met17*Δ on sulfate is mediated by hydrogen sulfide

Density-dependent growth in microbial populations is often mediated by the release of a chemical which, upon reaching a critical concentration, enables cells to grow and divide. Because *met17*Δ are perturbed at the enzymatic assimilation of the volatile compound hydrogen sulfide (H_2_S), we asked whether the growth mediator of *met17*Δ might be volatile. To test this, we limited gas exchange by using parafilm to seal the loose-fitting plastic lids on the culture tubes, and asked if the growth outcomes of *met17*Δ populations at different cell densities were impacted. Indeed, populations in parafilm-sealed tubes grew faster than in those without sealing (compare Fig 2B with Fig 2A; see S2 Fig for S288C). Even cultures at low cell densities eventually grew to saturation, although lag time exhibited stochasticity. When cultured in a 96-well plate sharing headspace, all cell densities could grow to saturation within a short duration (Fig 2C), suggesting that volatiles released from high-density cultures could facilitate the growth of low-density cultures. Interestingly, little stochasticity was observed in the lag time when lower cell densities grew in a shared headspace, suggesting that small differences amidst the gaseous environments within culture tubes gave rise to the stochastic growth dynamics (compare light gray trendlines in Fig 2C and Fig 2B). In contrast, only residual growth was observed when different initial cell densities of another methionine auxotroph, *met14*Δ, were grown on sulfate in a 96-well plate (Fig 2D, all cultures mildly increasing their turbidity by a fixed fold without reaching saturation).

**Figure 2.**
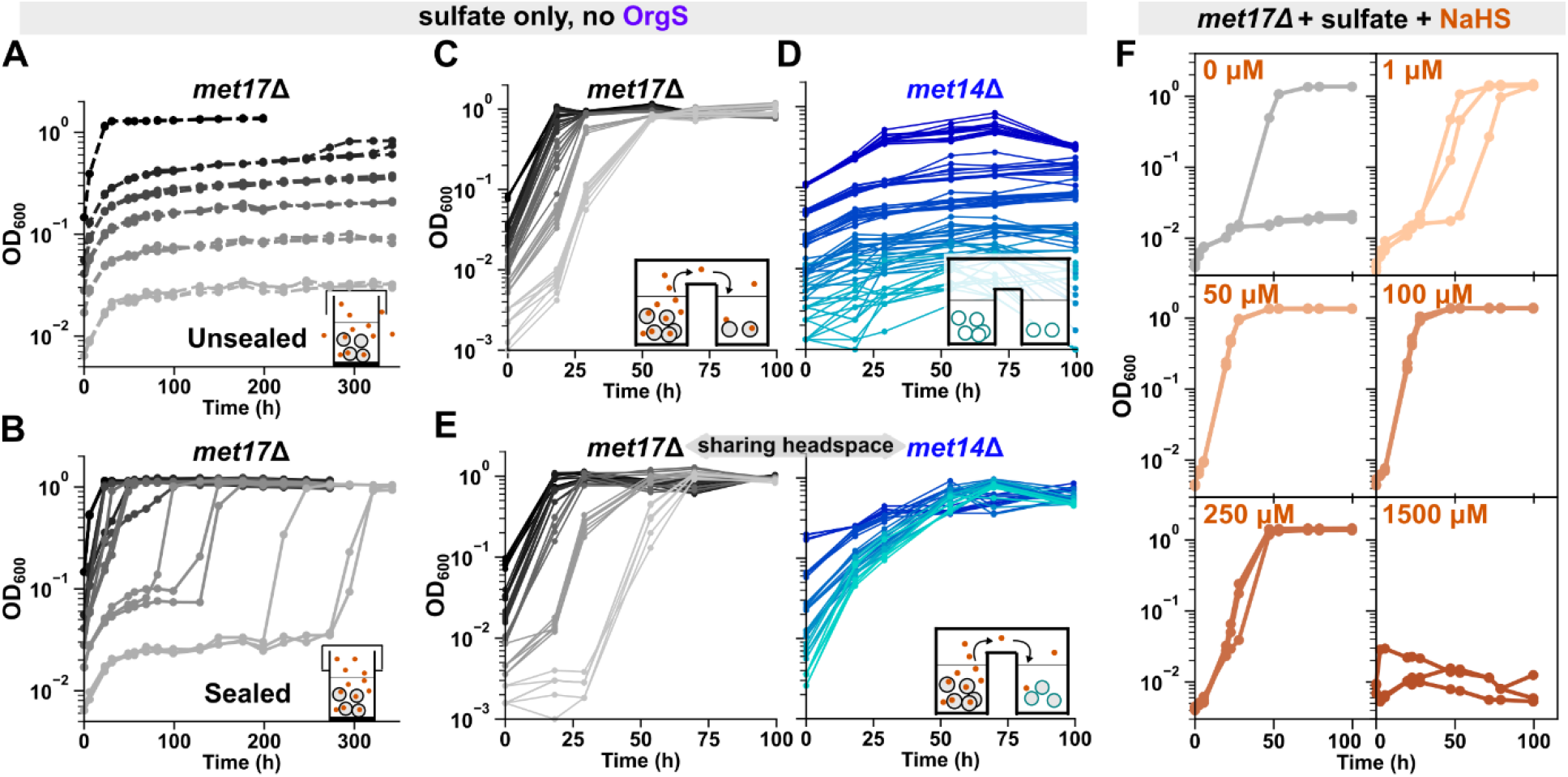
Hydrogen sulfide gas mediates the density-dependent growth of *met17*Δ on sulfate. **A, B**) Preventing gas escape allows low-density *met17*Δ cultures to grow. Growth curves of *met17*Δ yeast (WY2035) in minimal SD medium, cultured in glass tubes either covered only with plastic lids (**A**, “Unsealed”) or additionally sealed with parafilm around the plastic lids (**B**, “Sealed”). Darker shades of gray represent higher initial cell densities. Each trendline represents a 3-ml culture in a tube of 13-mm diameter. At any given initial cell density, six identical cultures were initiated, three each for the unsealed and sealed treatment. Note that even low-density cultures could eventually grow to saturation when gas exchange was limited by parafilming, indicating that the accumulation of a gaseous growth mediator was enabling population growth. **C**) Gas from high-density *met17*Δ cultures allows low-density *met17*Δ cultures in the same plate to grow on sulfate. Growth curves of *met17*Δ (WY2548) at different initial cell densities (indicated by shade of gray) growing in different wells of the same 96-well plate sealed with parafilm. All densities can grow deterministically when the headspace is shared in a multiwell plate. **D**) At all densities, *met14*Δ only showed residual growth in sulfate. Growth curves of *met14*Δ (WY2539) at different initial cell densities (indicated by shade of blue) growing in different wells of the same 96-well plate. **E**) Gas from *met17*Δ cultures allows *met14*Δ cultures to grow on sulfate. Growth curves of *met17*Δ (left panel, gray, WY2548) and *met14*Δ (right panel, blue, WY2539) at different initial cell densities (indicated by shades) sharing headspace in different wells of the same 96-well plate. **F**) Growth of low-density *met17*Δ on sulfate can be promoted by sodium hydrosulfide (NaHS), although high sulfide concentrations are toxic. At each sulfide concentration, each trendline represents one of three identical 3-ml liquid cultures (WY2531) in tubes sealed with plastic lids and parafilm. At 1500 μM, population growth was inhibited. All *met* mutants were in the RM11 background.

Corroborating previous reports of H_2_S release from *met17*Δ cells [15,26], we noted that *met17*Δ cultures growing on sulfate emanated a strong rotten-egg odor, and a piece of lead acetate paper placed in the headspace of *met17*Δ cultures turned black (S3 Fig). To test whether yeast can uptake and consume ambient H_2_S gas, we grew *met14*Δ yeast and *met17*Δ yeast in different wells of the same 96-well plate. Since *MET14* functions upstream of H_2_S formation during sulfate assimilation (Fig 1A), we hypothesized that *met14*Δ yeast should grow in the presence of H_2_S released by *met17*Δ. Indeed, in contrast to *met14*Δ showing only residual growth on sulfate by themselves (Fig 2D), all densities of *met14*Δ could grow when they shared headspace with *met17*Δ growing on sulfate in a 96-well plate (Fig 2E, right panel). Interestingly, growth of low-density *met17*Δ populations slowed down in the presence of *met14*Δ (compare Fig 2E left panel with Fig 2C), suggesting that the growth dynamics of *met17*Δ cultures were governed by the available quantity of the volatile sulfur forms which *met14*Δ and *met17*Δ competed for.

Finally, the growth of low-density *met17*Δ cultures was promoted by sodium salts of sulfide over a range of concentrations (Fig 2F and S4 Fig). Note that upon acidification, sulfide ions from the salts get protonated to release H_2_S gas. At a low sulfide concentration (1 μM), we see stochastic growth dynamics amidst three technical replicates (Fig 2F upper right panel). In contrast, the three replicates grew deterministically at intermediate concentrations of 50 μM and 100 μM (Fig 2F middle row). Thus, the transition from stochastic to deterministic growth dynamics observed with increasing initial cell densities in *met17*Δ cultures (Fig 1D) likely resulted from higher sulfide concentrations in high cell-density cultures. Interestingly, we noted that the cultures could not grow at the highest concentration of sodium hydroxide (NaHS) tested (1.5 mM in Fig 2F), indicating that high concentrations of sulfide are toxic to yeast cells, consistent with earlier studies [15,27]. This deterrence to the growth of eukaryotic cells might result from the well-characterized inhibition of mitochondrial cytochrome c oxidase by H_2_S [28].

Compared to RM11 *met17*Δ, S288C *met17*Δ required more H_2_S to grow and released less H_2_S. It took considerably higher concentrations of sodium sulfide to elicit growth in low-density cultures of S288C *met17*Δ on sulfate (S4 Fig, compare B with A). Moreover, S288C *met17*Δ could grow faster in the vicinity of RM11 *met17*Δ than when growing by themselves in a multi-well plate (S5 Fig, compare right panel of C with B), suggesting that RM11 *met17*Δ released more H_2_S than S288C *met17*Δ. Overall, both lower H_2_S release and higher H_2_S requirement may contribute to the sluggish growth of S288C *met17*Δ on sulfate compared to RM11 *met17*Δ (compare Fig1 with S1 Fig).

### *HSU1* is required for sulfide assimilation in *met17*Δ

The anomalous growth of *met17*Δ on sulfate indicates the existence of a *MET17*-independent pathway of sulfate assimilation in *S. cerevisiae*. Indeed, *met17*Δ could not grow at any cell density on sulfate-free medium (S6 Fig). During sulfate assimilation, Met17 catalyzes the reaction of sulfide with *O*-acetyl homoserine (OAH) to yield homocysteine (Fig 3A). *In vitro*, Met17 can also react sulfide with *O*-acetyl serine (OAS) to yield cysteine [29]. Other yeast species such as *K. lactis, Y. lipolytica* and *S. pombe* have dedicated enzymes to catalyze these two reactions [30]. Hypothetically, the *MET17*-independent pathway of sulfide fixation could meet the cell’s organosulfur requirements by catalyzing either of these reactions (*GENE X* in Fig 3A). However, alternate enzymes with similar functions have thus far not been characterized in *S. cerevisiae*. To discern if the alternative pathway requires OAH, we examined if *met2*Δ mutants, which lack OAH synthesis [31,32], could also bypass organosulfur auxotrophy. In a 96-well plate, where populations of *met2*Δ at different cell densities shared headspace, we found that the populations only showed residual growth (S7 Fig). Thus, it is likely that, similar to *MET17*, the alternative mechanism also utilizes OAH and sulfide.

**Figure 3.**
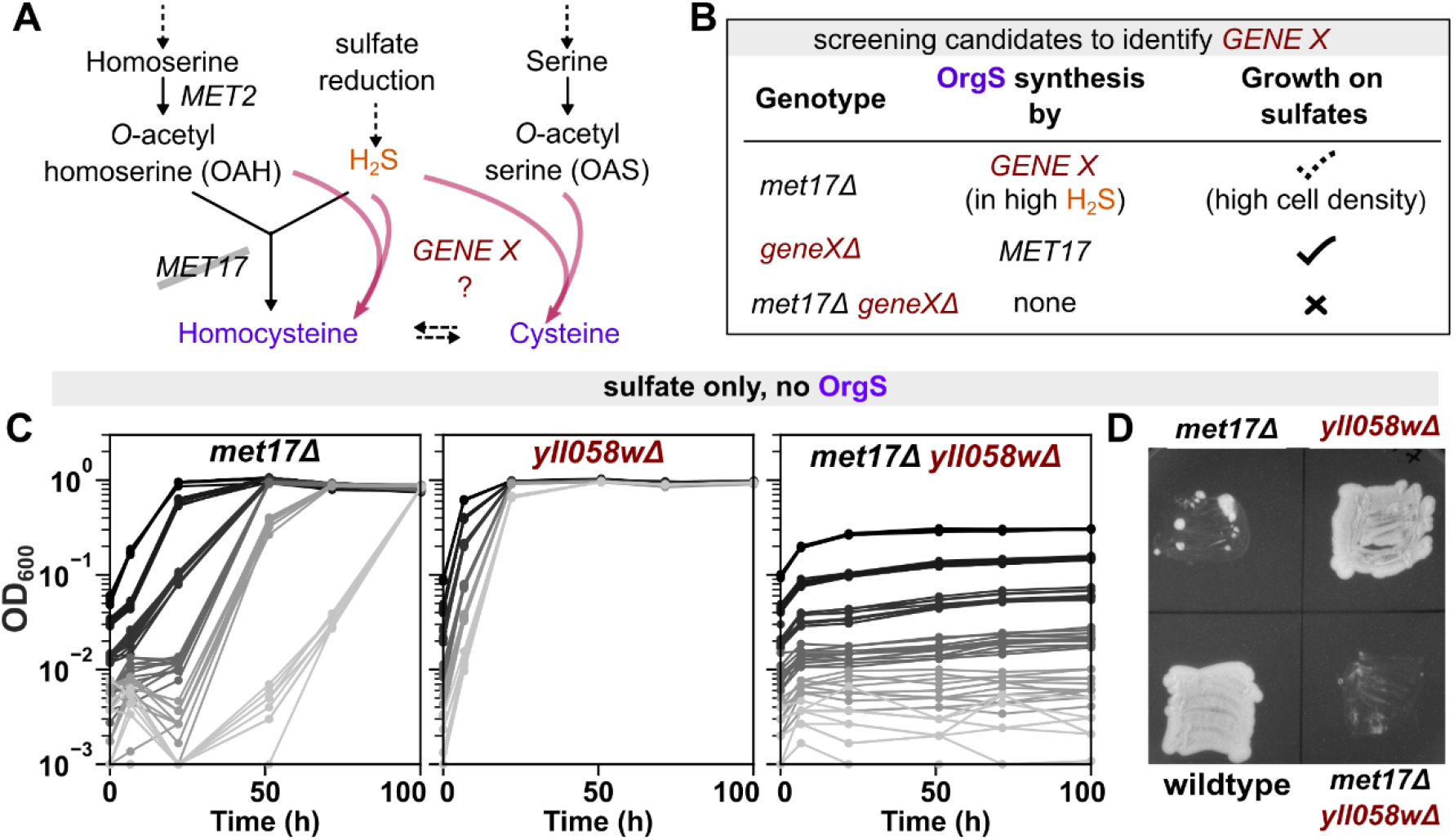
*HSU1* is required for the growth of *met17*Δ on sulfate. **A**) Schematic representing how an unknown *GENE X* could bypass the requirement for *MET17* by performing the enzymatic synthesis of either homocysteine or cysteine in a reaction where hydrogen sulfide is a substrate. Either of the two reactions would be sufficient to support the cell’s organosulfur requirement as organosulfurs (purple) are interconvertible. Solid and dashed arrows represent one-step and multi-step reactions, respectively. **B)** Schematic representing the genetic screen to identify *GENE X*. **C)** *YLL058W* is the hypothesized *GENE X*. Left panel shows that *met17*Δ of the S288C background (WY2590) can grow on sulfate at different initial cell densities when the populations share headspace in a 96-well plate. Middle panel shows that deletion of *yll058w* (WY2597) does not result in an auxotrophy, and all cell densities can grow without lags on sulfate in a 96-well plate. Right panel shows that double deletants of *met17* and *yll058w* (WY2595) can no longer grow on sulfate at any density in a 96-well plate. Altogether, these data indicate that *YLL058W* (or *HSU1*) is the *GENE X* hypothesized in panel **A. D**) Unlike *met17*Δ (WY2590), the double mutants *met17Δyll058w*Δ (WY2595) do not show papillae when patched onto agar plates containing SD minimal medium. *Yll058w*Δ (WY2597) grow dense lawns similar to wildtype yeast (WY2601). All four genotypes were patched onto the same agar plate which was sealed with parafilm and imaged after 5 days.

Met17 is a pyridoxal phosphate–dependent enzyme. We proposed that any protein that performs a catalytic function similar to Met17 will bear structural similarity to the Met17 enzyme. Five candidate genes were identified by a protein BLAST using Met17’s sequence: *CYS3, STR3, ICR7, YLL058W, YHR112C*. If any of these genes was *GENE X*, then the *met17Δ geneX*Δ double mutant would no longer grow on sulfate even at a high initial cell density (Fig 3B). However, *geneX*Δ single mutant should grow on sulfate due to the activity of *MET17* (Fig 3B). Out of the candidates, *cys3*Δ did not meet this criterion of our screen since it is a known cysteine auxotroph [33]. As a quick method of screening the remaining four candidates, we leveraged the fact that *met17*Δ is a background mutation in the yeast deletion library derived from the strain BY4741 of the S288C background [21]. For each candidate, different initial cell densities of the same mutant were inoculated into different wells of the same 96-well plate. This set-up allows us to clearly distinguish between residual growth (e.g., *met14*Δ, Fig 2D) and growth to saturation (e.g. *met17*Δ, Fig 2C). Out of the candidates tested, only deletion of *YLL058W* abrogated growth of *met17*Δ on sulfates (S8 Fig).

To confirm that the *yll058w*Δ single mutant is not auxotrophic, we deleted the gene in the S288C strain background. Indeed, while *met17*Δ grew in a density-dependent fashion (Fig 3C left panel), *yll058w*Δ yeast could readily grow on sulfate without supplementation of organosulfurs (Fig 3C middle panel). In contrast, double mutants of *met17Δyll058w*Δ could no longer grow to saturation on sulfate at any density (Fig 3C right panel, showing only residual growth). The double mutants did not show papillae even when patched onto the same solid SD medium plate as H_2_S-releasing *met17*Δ (Fig 3D; S9A Fig for RM11), and could not grow even when cultured in the same 96-well plate as *met17*Δ (S9B Fig), indicating that the inability of *met17Δyll058w*Δ to grow on sulfate did not result from impairments in sulfide release but from defects in sulfide assimilation. While we were investigating this phenomenon, two other groups simultaneously reported this sulfide-assimilation function of *YLL058W* [24,25], and Yu *et*.*al*. named the locus *Hydrogen Sulfide Utilizing-1* (*HSU1)* [25]. We will henceforth use the name *HSU1* to refer to *YLL058W*. These other groups additionally demonstrated that Hsu1 protein can catalyze the same biochemical reaction as *MET17*, albeit at considerably lower efficiency [24,25].

### *HSU1* may have diverse roles in sulfur metabolism

Based on the fact that *HSU1* can assimilate sulfide when sulfide concentrations are high, we hypothesized two possible functions for *HSU1*: 1) an alternate pathway to maximize sulfur assimilation when cells experience sulfur starvation; and/or 2) a mechanism to neutralize excess sulfide when ambient sulfide concentrations get dangerously high (e.g. Fig 2F last panel). Correspondingly, gene expression might be triggered either by sulfur starvation or by exposure to high sulfide concentrations. To test these hypotheses, we introduced a GFP coding sequence before the stop codon on the C-terminus of *HSU1* in the S288C and the RM11 background. We then imaged these cells after treatment with sulfate-free medium (sulfur starvation; Figs 4A and 4C) or after exposure to different concentrations of sodium sulfide (Figs 4B and 4D). In both backgrounds, exponentially growing wildtype cells showed little GFP signal (Figs 4A and 4C, left panels). Interestingly, a sharp increase in GFP fluorescence was observed under sulfur starvation (Figs 4A and 4C, right panels). Non-specific starvation may not induce Hsu1, as cells grown to stationary phase in the SD minimal medium have little Hsu1 protein (Figs 4B and 4D, left panels). When minimal medium is supplemented with an intermediate level of sulfide (0.6 mM), Hsu1 protein is induced—more so in the RM11 than in the S288C background (Fig 4B and 4D, middle panels). At a high sulfide concentration (1.8 mM which can inhibit the growth of wildtype cells (S12A Fig), Hsu1 protein is induced in both backgrounds. Hsu1 protein generally shows a diffused localization in the cytoplasm, although punctate localization was occasionally also observed under sulfide exposure (Fig 4B and 4D, middle and right panels).

**Figure 4.**
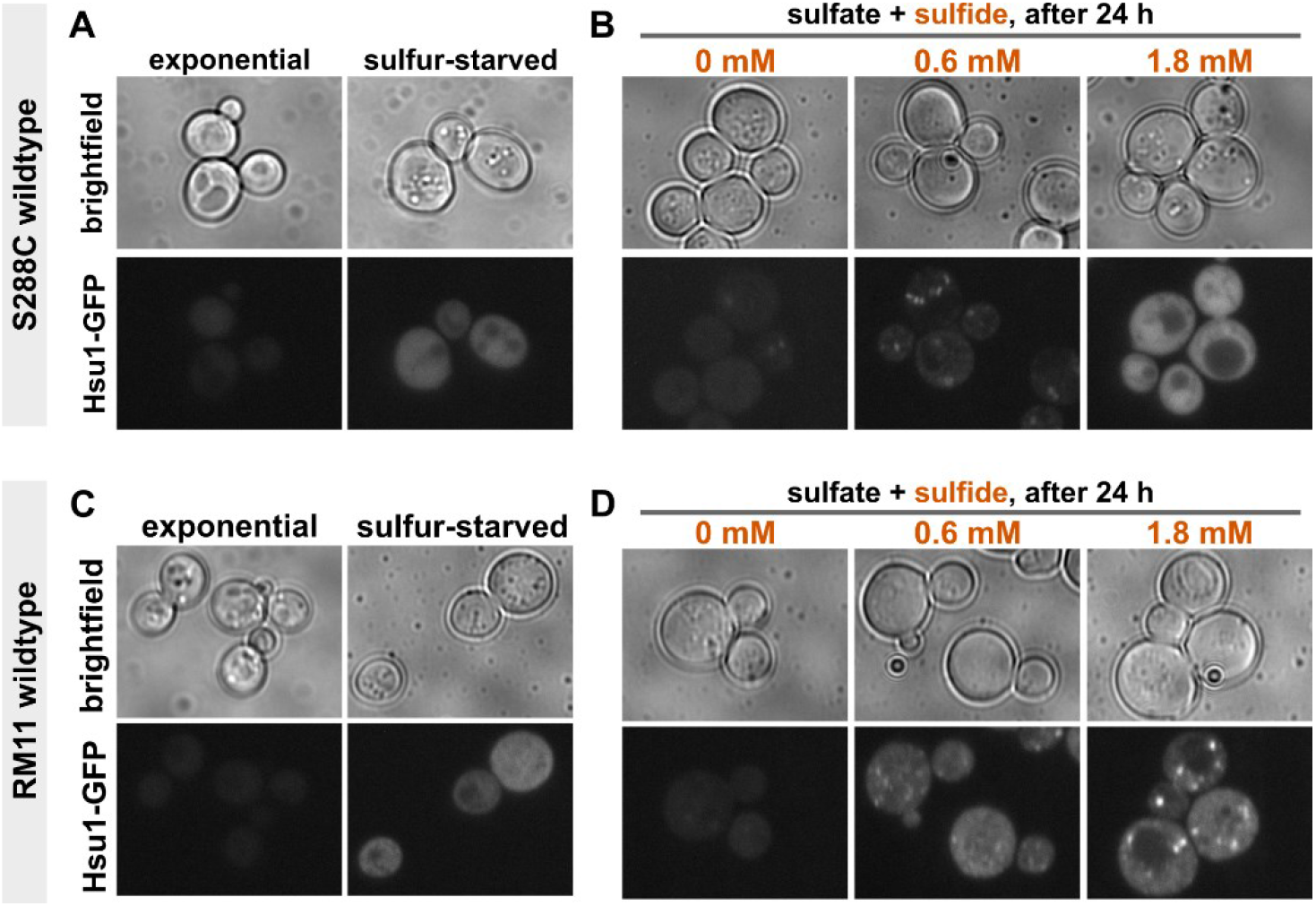
*HSU1* may have diverse roles in the sulfur metabolism of wildtype yeast. The expression profile of Hsu1-GFP in wildtype yeast of S288C (WY2616; **A-B**) and RM11 (WY2618; **C-D**) backgrounds suggest multiple functions of *HSU1* in sulfur metabolism. **A, C** Sulfur starvation induces expression of Hsu1-GFP in wildtype yeast within 4 hours. Hsu1-GFP is not detected in exponential phase yeast growing in SD minimal medium (left columns in **A** and **C**). **B, D** Adding sulfide (Na_2_S) to SD minimal medium also induces expression of Hsu1-GFP in wildtype yeast. Localization of Hsu1 is largely diffused throughout the cytoplasm, although puncta can be observed under some conditions of sulfide exposure. Note that 1.8 mM sulfide impairs growth of wildtype yeast (S12A Fig), but cells largely remain viable after 1 day of treatment (as judged by flow cytometry and plating in S12C Fig). No fitness disadvantage was detected for *hsu1*Δ in comparison to wildtype yeast in a range of sulfur environments (Figs S11-S13).

To identify the functional significance of *HSU1*, we compared wildtype and *hsu1*Δ S288C yeast under various conditions, including during sulfur starvation or sulfide exposure. Consistent with the lack of *HSU1* expression in exponential phase, the growth rate of *hsu1*Δ did not differ from wildtype during exponential growth on minimal medium (S10 Fig). In competition assays under sulfur starvation, variation in the outcome was high, and no trend was detected in the fitness difference between wildtype and *hsu1*Δ (S11 Fig). High sulfide concentrations did impair growth, as was previously reported for *met17*Δ (Fig 2F), but the responses of wildtype and *hsu1*Δ to high sulfide were very similar (S12 Fig). Thus, *HSU1* does not provide any discernable advantage to wildtype yeast under either sulfur starvation or sulfide exposure.

Recently, quantitative trait locus mapping of wine strains revealed that polymorphisms in *HSU1* affect the production of volatile dimethyl sulfide from the exogenous organosulfur S-methylmethionine (SMM) [34]. Yeast usually obtain SMM from plant sources (e.g. from grape juice), and can grow on SMM as the sole sulfur source [35]. Interestingly, *HSU1* resides on the same chromosomal segment as the SMM permease *MMP1* and the SMM-metabolizing enzyme *MHT1* [36], and all three enzymes are transcriptionally upregulated by Met4 [37], which in turn is required for growth on SMM [36]. To test if *HSU1* contributes to SMM metabolism, we compared the growth rate of wildtype and *hsu1*Δ yeast in medium containing SMM as the sole sulfur source. However, both genotypes had indistinguishable growth rates (S13 Fig). Thus, while Hsu1 expression suggests multiple roles of the protein in sulfur metabolism, the precise function of *HSU1* in wildtype yeast remains unclear.

### A mathematical model assuming low efficiency of sulfide assimilation can explain density-dependent growth of *met17*Δ

What mechanisms might give rise to the density-dependent growth of *met17*Δ on sulfate? One possibility is stochastic cell-state switching, which has been widely observed in diverse microbes [38]. Specifically, in the case of *met17*Δ (Fig 5A), all cells may start out being “inactive”– incapable of assimilating sulfide for growth. However, the accumulation of sufficient sulfide may trigger a switch to an “active” or growth-competent state where cells can fix sulfide and give birth to new cells, which in turn release more sulfide. A sulfide-dependent regulation of *HSU1*’s expression may serve as the mechanism for such a switch. If this were the case, we would expect Hsu1 protein to be absent or weakly expressed in cells of low-density *met17*Δ cultures that fail to grow. However, considerable Hsu1-GFP expression was observed in low-density cultures within 4 hours of transfer to minimal medium, even if the cultures did not eventually grow (Fig 5B). *HSU1* expression in *met17*Δ probably occurs because organosulfur starvation in *met17*Δ leads to a similar molecular response as sulfur starvation in wildtype cells [39], and we have shown that Hsu1 is induced in the latter condition (Fig 4A,C). Thus, if a cell-state switch for sulfide assimilation exists, it is not modulated at the level of Hsu1 protein expression.

**Figure 5.**
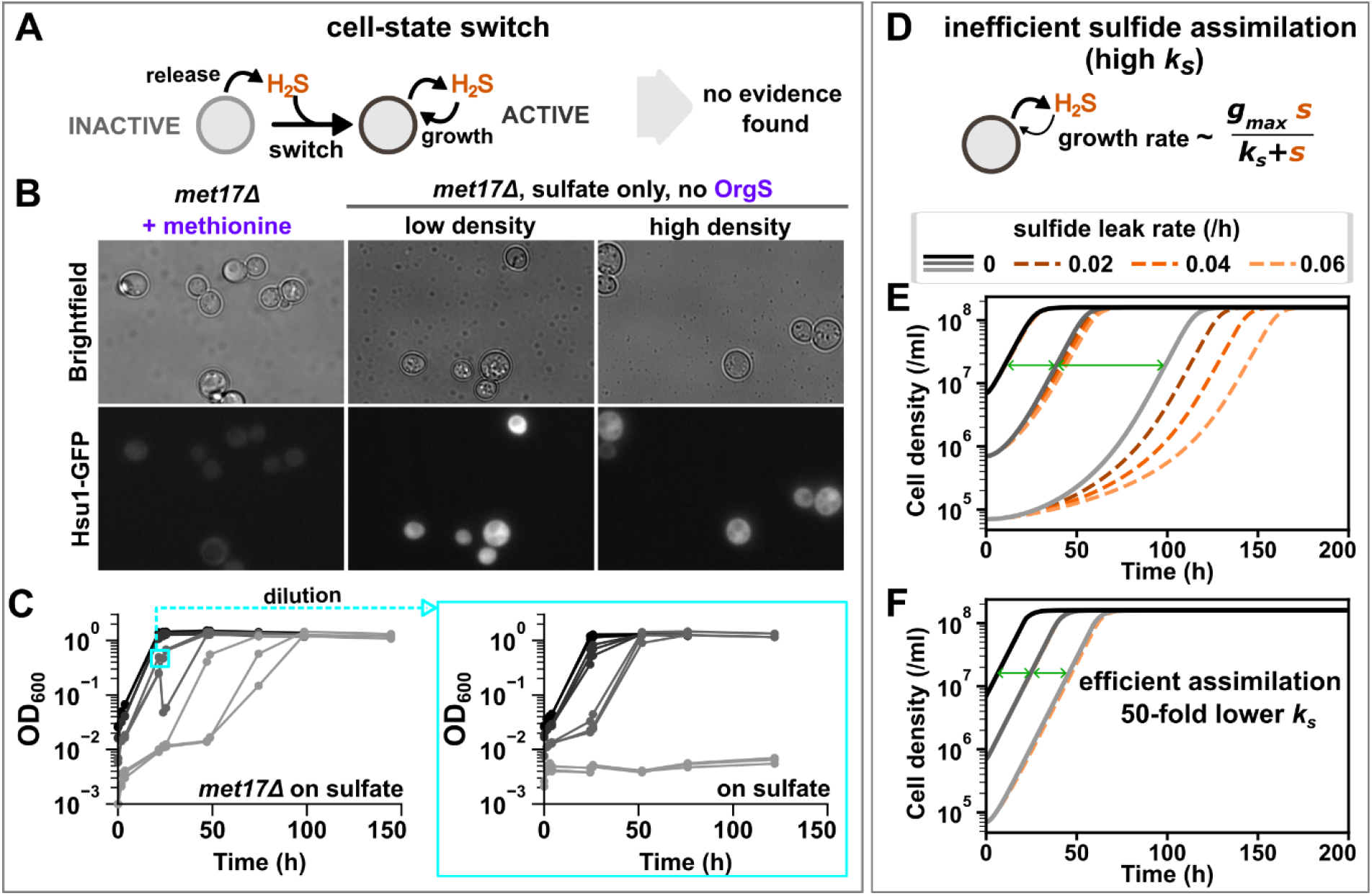
Inefficient sulfide assimilation can lead to density-dependent growth of *met17*Δ. **A**) One possible mechanism for density-dependent growth is that *met17*Δ cells switch to a growth-competent (sulfide-assimilating) state upon sufficient sulfide accumulation. **B**) Expression of *HSU1* cannot serve as the mechanism of a cell-state switch. While *met17*Δ (WY2623) growing exponentially in methionine-supplemented medium do not show much expression of Hsu1 protein, cells in both low-density and high-density cultures show expression within 4 hours of transfer to minimal medium (without methionine or other organosulfur supplements). **C**) *met17*Δ cells (WY2548) that have started to grow on sulfate continue to show density-dependence when diluted to fresh minimal medium. This suggests that there is no switch to a growth-competent state in *met17*Δ cells or that the switch is very transient and does not persist dilution. Lighter gray shades indicate lower initial cell densities. Each trendline represents a 2.5-ml culture in a parafilm-sealed 13-mm glass tube. Note that since experiments are initiated with exponentially growing cells, longer lags at lower cell densities are the hallmark of density-dependent growth. **D-F**) Inefficient sulfide assimilation by Hsu1 can give rise to density-dependent growth. **D**) All *met17*Δ cells are growth-competent, with the key assumption being that cell growth rate has a Monod-dependence on sulfide concentration. **E**) A mathematical model (see main text) can generate density-dependent growth dynamics using parameters measured in or inferred from experiments. Lags (green double arrows) are longer at lower cell densities (lighter gray shades). Lag times at lower cell densities are also sensitive to loss or leakage of sulfide (dashed lines). **F**) Both longer lags and the sensitivity to sulfide loss disappear when the efficiency of sulfide assimilation is improved. The sulfide concentration of half-maximal growth rate (*k*_*s*_) was lowered by 50-fold to simulate better sulfide assimilation. Initial cell densities in simulations for **E-F** were equivalent to an OD_600_ of 0.001 (light gray), 0.01 (gray) and 0.1 (black), respectively.

A cell-state switch could however operate by other molecular mechanisms. For instance, a post-translational modification might activate Hsu1 once the ambient sulfide concentration has become sufficiently high. Independent of molecular mechanisms, we tested the existence of a cell-state switch by taking cells from a *met17*Δ culture that had started to grow on sulfate and using them to re-initiate cultures at different initial cell densities in minimal medium (containing sulfate). We hypothesized that since Hsu1 has already been activated, the fresh inoculations would be able to grow at lower initial densities. However, this was not observed (Fig 5C), indicating that there is no switch to a growth-competent state in *met17*Δ cells or that the switch does not persist dilution. Thus, a H_2_S-dependent growth switch is unlikely to be the sole mechanism producing density-dependent growth of *met17*Δ on sulfate. Similar results were observed for *met17*Δ of S288C background (S14A Fig).

Alternatively, density-dependence could simply result from inefficient assimilation of sulfide by Hsu1. To test this hypothesis, we developed a mathematical model describing how *met17*Δ grow by releasing and consuming sulfide. Note that we were able to use a deterministic model instead of a stochastic one because our data suggested that the observed stochastic growth dynamics resulted from experimental factors, rather than biological ones: Low-density *met17*Δ populations only show stochastic lag times in individual tubes (Fig 2B), and not when they share headspace in a multi-well plate (Fig 2C). This suggests that the variance in lag times results from small differences in the gas environments that the cells experience in each tube, which in turn could result from minor defects in sealing.

The model, comprising two differential equations describing the dynamics of population density *x* and sulfide concentration *s*, assumes that (i) all cells are capable of growth, with a carrying capacity *K*, (ii) growth rate has a Monod-dependence on sulfide concentration (Fig 5D), with maximal growth rate *g*_*max*_ and Monod constant (sulfide concentration at which half-maximal growth rate is achieved) *k*_*s*_; and (iii) sulfide is produced by cells (via sulfate reduction) at a constant rate *r*, consumed by cells at *c* amount per birth, and lost at a rate of *δ*. The equations are as follows:

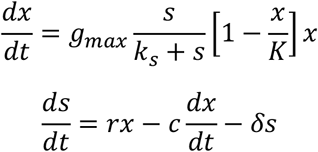

Some parameters were directly measured in experiments: *c* = 3 fmole/cell (S15 Fig), *K* = 1.6 × 10^8^ cells/ml; *g*_*max*_ = 0.26 /h (Methods: “Mathematical model”). Others were inferred by fitting the model to the experimental data: *r* = 0.41 fmole/cell/h and *k*_*s*_ = 7.1 μM (Supplementary Text 1, section 3). This *k*_*s*_ includes a factor that accounts for liquid-gas portioning of sulfide (Supplementary Text 1, section 4).

When population density is much smaller than the carrying capacity, growth rate will have a Monod-dependence on sulfide concentration. While this relationship is similar in form to the Michaelis-Menten enzyme kinetics equation, the two equations are not equivalent. Michaelis constant and Monod constant respectively describe the substrate concentration at which half-maximal enzyme activity and half-maximal cell growth is attained. Cell growth is not only a function of enzyme affinity for substrate, but also of additional factors such as the number of enzymes in the cell and the coordination between the reaction and the rest of cellular metabolism. Indeed, Van Oss et al. report that the Michaelis constants of both Hsu1 and Met17 for sulfide are in millimolar range [24], but that concentration would be toxic to yeast (Fig 2F and S12A Fig). Thus, parameters measured in *in vitro* enzyme assays of Hsu1 cannot be applied here.

This model reproduced the density-dependent growth of *met17*Δ on sulfate: the lag times at lower densities are longer than those at higher densities. Specifically, for three cultures separated by equal (10)-fold differences in initial densities, exponential growth curves will have an equal temporal distance in a semi-log plot. Yet for *met17*Δ, the spacing between the lower-density curves is longer than that between the higher-density curves (Fig 5E, green double arrows; compare with Fig 2B). Interestingly in our model, low-density cultures are more sensitive to loss of sulfide (Fig 5E), potentially explaining the stochastic lag times observed in culture tubes (Figs 1D and 2B). In contrast, small variations in initial cell densities, such as those attributable to pipetting error, had a negligible effect on growth dynamics (Supplementary Text 1, section 3), presumably because the lowest cell density in our experiments was already quite large (∼10^5^ cells/ml). Finally, the efficiency of sulfide utilization impacts density-dependent growth: lowering the Monod constant *k*_*s*_ by 50-fold not only abrogated lag at all cell densities, but also the sensitivity to sulfide loss (Fig 5F). Thus, low efficiency of sulfide assimilation may be sufficient to explain the density-dependent growth we observe for *met17*Δ on sulfate.

## DISCUSSION

In this study, we have investigated how budding yeast cooperatively overcome the metabolic defect caused by the deletion of *MET17*. The enzyme Met17 was believed to be required for yeast to generate organosulfurs from inorganic sulfate [23,40]. However, we have shown that *met17*Δ is not a true organosulfur auxotroph and can in fact grow using sulfate as the sole sulfur source in a density-dependent manner. This phenomenon occurs because the locus *YLL058W/HSU1* can take over Met17’s function of organosulfur synthesis, albeit only at relatively high sulfide concentrations. Specifically, after reducing sulfate to sulfide, *met17*Δ cells fail to react sulfide with OAH to form homocysteine. The accumulating sulfide readily crosses cell membranes and partitions between the liquid and air phases. While *HSU1* is expressed in *met17*Δ experiencing organosulfur starvation (Figs 5B and S14A), it is likely that Hsu1 is inefficient at sulfide assimilation [24,25]. Therefore, ambient sulfide levels need to be considerably high before Hsu1 can synthesize sufficient organosulfurs to fuel cell division. Cultures starting at low cell densities take longer to reach a sufficient level of sulfide, or may not reach it at all as sulfide can be lost due to gas exchange or oxidation. Thus, depending on the starting cell density and other experimental factors (e.g. the extent of aeration), *met17*Δ can either grow deterministically, stochastically or not grow at all on sulfate (Figs 1 and 2). The stochastic growth dynamics of low-density *met17*Δ culture is unlikely to be caused by a sulfide-induced cell-state switch (Fig 5A-C). Instead, low activity of Hsu1 [24,25] results in density-dependent growth (Fig 5D-F). Furthermore, at low initial cell densities, growth is particularly sensitive to variations in ambient H_2_S gas levels (Fig 5E), which can explain the stochasticity observed in the growth outcome of *met17*Δ on sulfate.

### Reconciling discrepancies in experiments in yeast genetics and metabolism

In retrospect, the erratic growth behaviour of *met17* mutants in the absence of organosulfurs had been noticed prior to our study. Even though *MET17, MET15* and *MET25* were identified as the same locus [13–15], mutants of some *met25* and *met17* alleles failed to grow unless supplemented with organosulfurs [14,23], whereas mutants of other *met17* alleles could grow when supplied with sulfite or sulfide [23]. A careful inspection reveals that even in the study where Cost and Boeke proposed *met15* as a selection marker, *met15* mutants exhibited some low level of growth when patched onto sulfate-containing minimal medium which lacks organosulfurs [41]. Finally, when selecting for methyl mercury resistance, *met15* (and *met2*) mutants grew despite the medium not having any organosulfurs [42,43]. While some of these observations could be due to reduced-function mutations associated with specific alleles, it is likely that the leaky auxotrophy of *met17* mutants that we have elucidated in our study could explain some of these perplexing observations.

During the course of this study, two other studies were published reporting the anomalous growth of *met17*Δ and identifying the locus *YLL058W/HSU1* as the cause [24,25]. However, the two studies disagreed on the observed effect of sulfide on *met17*Δ. Van Oss *et al*. claimed that sulfide accumulation was toxic to *met17*Δ cells, and therefore, *met17*Δ could grow only once *HSU1* became active and neutralized some of the sulfide [24]. In stark contrast, Yu *et al*. claimed that sulfide accumulation facilitated *HSU1*-dependent growth of *met17*Δ [25]. Consistent with Yu *et al*., we found that a range of sulfide concentrations can promote the growth of both S288C *met17*Δ and RM11 *met17*Δ on sulfate (Fig 2F; S4 Fig). Consistent with Van Oss *et al*., we found that high sulfide concentrations can be toxic to yeast cells but we only observe such growth inhibition with addition of extraneous sulfide (Fig 2F; S4 Fig; S12 Fig). In addition, facilitating gas accumulation by sealing the tubes promoted the growth-propensity of *met17*Δ (Fig 2A-B), indicating that *met17*Δ usually experienced favorable sulfide regimes in our experiments. We can think of two possible explanations for the disparate results in Van Oss *et al*. First, the growth-promotion effect of the sulfide chelator Fe(III)-EDTA may not result from a reduction in sulfide levels. For instance, instead of removing sulfide, the chelator might have improved bioavailability by concentrating the sulfide, or growth might have been promoted simply by Fe(III)-supplementation. Either of these mechanisms would explain why prototrophs in their study showed an improved growth in the presence of the chelator (Fig 4B in [24]). Future studies could address this possibility by using less invasive techniques of sulfide removal such as suspending lead acetate paper in the headspace. Second, despite being closely related, the FY4 strain used in Van Oss *et al*. could differ from the S288C strain used in Yu *et al*. (and our study) in sulfide release rate or in sensitivity to sulfide.

Contrary to our findings of no detectable fitness effects of *hsu1*Δ compared to wildtype either during sulfur starvation or exposure to toxic Na_2_S concentrations (S11A Fig; S12), Yu *et al*. detected a small defect for *hsu1*Δ under sulfur limitation and a small advantage for *hsu1*Δ in abundant sulfur [25]. However, Yu *et al*. used colony counting, and we noted that colony counting produces data with high variance (S11 Fig). This variance arises from the stochasticity associated with plating small numbers of colonies, as well as errors in counting colonies (either manually or using automation). This makes it difficult to detect small fitness differences through colony counting without a high number of replicates. Our analysis, which is based on flow cytometric counting of tens of thousands of cells, has considerably higher precision.

Overall, our work has implications for yeast genetics experiments. *met17*Δ is a commonly used selection marker, featuring as a background deletion in the yeast deletion collection [21,44]. An understanding of the mechanism of its leaky auxotrophy will allow researchers to design protocols that circumvent the caveat. For instance, *met17*Δ can still be used as a selection marker if patching is done at low cell densities and growth phenotypes are assessed within short time intervals.

### Considerations for quantitatively investigating volatile-mediated microbial interactions

The volatile nature of the growth substrate H_2_S can lend high variability to experimental observations. Our work raises awareness towards two factors in particular: First, volatile nutrients can be exchanged between seemingly unconnected microbial populations. By showing that *met14*Δ yeast can grow in the vicinity of sulfide-releasing *met17*Δ (Fig 2E), we demonstrate that assessing auxotrophy in a set-up where multiple mutants share headspace can lead to confounding conclusions. For example, H_2_S from neighbouring unsealed plates could explain the surprising repeated growth of wildtype FY4 in the absence of inorganic or organic sulfur observed by Van Oss *et al*. (Fig 2D of [24]). Second, small differences in experimental setups could give rise to considerably different experimental outcomes. A mathematical model describing the growth of *met17*Δ on sulfate reveals that, at lower cell densities, growth outcome is sensitive to loss of sulfide (Fig 5E). While loss of non-volatile substrates primarily occurs through chemical degradation, sulfide could also be lost due to gas escape from the culture chamber, which in turn is affected by experimental factors such as temperature [25], the thoroughness of tube sealing, and the use of agitation during growth. The problem is further exacerbated by the non-monotic effect of sulfide on *met17*Δ yeast populations: growth-promotion at lower doses and growth-inhibition at higher doses. Growth outcomes are thus sensitive to the exact sulfide environment that the yeast experience, which is difficult to quantitatively compare between different studies.

The liquid-gas partitioning of sulfide could pose an additional challenge for quantitatively comparing results from different setups. However, this hindrance may not be as severe as one would naively imagine. If the timescale of liquid-gas partitioning is much faster than the biological processes (e.g. consumption and birth) and gas leakage, then partitioning does not need to be modeled explicitly (Supplementary Text 1, section 4). Rather, partitioning can be accounted for by including a liquid-gas partition factor in the Monod constant (*k*_*s*_) and the leakage rate (*δ*).

### Possible functions of *HSU1* in sulfur metabolism and stress response

What might be the functions of *HSU1*? We could not detect any fitness advantage that *HSU1* confers to wildtype yeast during sulfur starvation, sulfide exposure or while utilizing SMM as the sole sulfur source (Figs S11-S13). However, two lines of evidence suggest that Hsu1 plays a role in sulfur metabolism. First, *HSU1* transcription is induced during sulfur limitation and starvation [37,45,46]. Hsu1 protein is also induced during sulfur starvation (Fig 4A, C) or limitation [25]. Second, *in vitro* biochemical assays show that, similar to Met17, Hsu1 can act as a homocysteine synthase (OAH sulfhydrylase) [24,25] as well as a cysteine synthetase (OAS sulfhydrylase) [25], although Hsu1 is much less active than Met17 [24,25].

*In vitro* assessment might however not suffice to reveal *HSU1*’s function since sulfur metabolism enzymes can show a high degree of substrate promiscuity (e.g. [47,48]). Given *HSU1*’s role in production of dimethyl sulfide from SMM, one can speculate that *HSU1* is involved in the production of volatile sulfur compounds. Such metabolites might have non-essential roles in cellular metabolism or confer a fitness advantage only in a specific environmental context. In line with this hypothesis, the *HSU1* gene shows considerable variation within the yeast strains represented in the *S. cerevisiae* Genome Database [49]. Of the 41 strains, *HSU1* contains large deletions in 5 strains, in addition to multiple polymorphic regions (S16 Fig). In contrast, *MET17* is highly conserved in all except one of these strains. This variation in *HSU1* does not result purely from its proximity to the telomere since genes such as *GTT2* are more proximal to the telomere but are still highly conserved. Thus, it is likely that *HSU1* plays a biosynthetic role in secondary metabolism which can tolerate more variation within yeast strains.

Alternatively or in addition, *HSU1* might function in stress response, which is connected to sulfur metabolism through glutathione [2]. Various findings support this speculation: The heat shock factor, which responds to diverse stresses, constitutively binds to the promoter region shared by *HSU1* and *GTT2* (glutathione S-transferase), and the binding is increased upon heat shock [50]. *HSU1* transcription is induced after heat shock, or by exposure to hydrogen peroxide or a strong reducing agent [51]. Upon exposure to heavy metal cadmium ion, cells induce transcription of sulfur-depleted alternatives of their abundant proteins and almost all genes involved in sulfur metabolism [52,53], including *HSU1* [52]. We found that in wildtype cells, Hsu1 protein level is induced upon exposure to high sulfide concentrations (Fig 4B, D) [25]. Yeast can release H_2_S during wine fermentation [54], either when nitrogen becomes limited in grape juice, or when elemental sulfur (from fungicides etc.) is reduced to H_2_S [55]. Thus, *HSU1* might function to neutralize high sulfide. Note that although Hsu1 localized throughout the cytoplasm, punctate localization was occasionally observed (depending on sulfide concentration and strain background; Figs 4B and 4D). Future studies could investigate if Hsu1 puncta colocalize with stress granules or with foci formed by other sulfur metabolism enzymes (e.g. Sam1 and Sam2 [56]).

In sum, our work provides important considerations for experimental design in both yeast genetics and volatile-mediated microbial interactions, and draws attention to less understood aspects of sulfur metabolism.

## METHODS

### Media, chemicals and culturing conditions

Prior to an experiment, strains were revived from glycerol stocks stored at -80ºC by streaking on YPD plates (2% agar included in liquid YPD: 10 g/L yeast extract, 20 g/L peptone, 20 g/L glucose) and incubating at 30ºC for 48 hours. 2 ml of liquid YPD was then inoculated from a single colony from the plate and grown overnight at 30ºC with agitation. These saturated YPD cultures were used to inoculate SD minimal medium (6.7 g/L Difco Yeast Nitrogen Base without amino acids but with ammonium sulfate from Thermo Fisher Scientific, Waltham, MA, USA and 20 g/L D-glucose) supplemented with any amino acids required by the auxotroph used in that experiment. For experiments with *met17*Δ, 20 mg/l of methionine was added. For WY2035, 20 mg/l of methionine and 30 mg/l lysine were added. For strains from the yeast deletion library used in Figs S5 and S8, 20 mg/l methionine, 20 mg/l uracil, 20 mg/l histidine and 60 mg/l leucine were added for exponential growth [57]. Sulfur starvation medium was prepared by replacing all sulfate salts in SD medium; a detailed list of ingredients and preparation notes are provided in S3 Table.

In certain experiments, minimal medium was supplemented with sodium sulfide, either NaHS or Na_2_S. For Figs 2F and S15, a 1 M NaHS stock solution was used (gifted by Prof. Mark Roth, FHCRC, Seattle, WA, USA). For all other experiments, a 1 M stock solution of Na_2_S (Fisher Scientific catalogue no. 10656811) was prepared in 0.001 M NaOH in an anaerobic chamber. Both the stock solutions were stored in an anoxic environment in glass tubes with self-sealing septa. Immediately prior to an experiment, approximately 50 μl of the sulfide stock was drawn out using a 27G-1_1/4_ gauge needle. The necessary amount was pipetted into the cell culture tubes on the glass wall and mixed by vortexing only after the tubes were sealed, to minimize loss of gaseous H_2_S during handling.

For S13 Fig, S-methylmethionine (catalogue no. M0644, Tokyo Chemical Industry Co Ltd., Tokyo, Japan) was added at a final concentration of 0.2 mM to sulfur starvation medium.

Glass tubes of either 13-mm or 18-mm diameter, with loosely fitted plastic lids, were used for culturing. Culture volume was maintained at either 2.5 ml or 3 ml in the smaller tubes or 7 ml in the larger tubes. Where sealing is mentioned, the tubes were additionally sealed with a layer of cling film and two rounds of parafilm. Tubes were placed either in a 30ºC incubator on a tilted rack with 250 rpm agitation or on a rotor wheel in a 30ºC warm room.

### Yeast strains

All yeast strains used in this study are listed in S1 Table. Yeast nomenclature follows the standard convention. Primers used in the study are listed in S2 Table.

Deletion strains were constructed either via yeast crosses or by homologous recombination [58,59]. Crosses were carried out by mating parent strains, pulling diploids, sporulation, tetrad dissection, and selection on suitable plates. As an example, *hsu1*Δ (WY2597) was constructed by PCR-amplifying the KanMX resistance gene from a plasmid (WSB26 [60]) using the primers WSO705 and WSO706, with a 45-base pair homology to the upstream and downstream region of the *HSU1* gene, respectively. A diploid heterozygous for *met17*Δ was then transformed with the PCR product and transformants were selected on a G418 plate. Successful deletion was confirmed via a checking PCR with a primer upstream of the *HSU1* gene (WSO707) paired with an internal primer for the amplified KanMX cassette (WSO144). This diploid was then sporulated, tetrads dissected, and replica plated onto G418 and minimal medium plates for genotyping, such that both *hsu1*Δ and *met17Δhsu1*Δ strains could be generated from this protocol. Mating-type genotyping of selected clones was carried out using a PCR-based protocol with primers WSO690-692 [61]. One exception to this standard deletion process was the generation of a marker-free deletion of *MET17* (WY2548) using the counter-selectable marker amdSYM [62].

For generating strains with different fluorescent labels for competition assays, an S288C MATα *met17Δhsu1*Δ strain (WY2602) was crossed with wildtype strains expressing either eGFP (WY1364) or mOrange (WY1376). The fluorophores tagged the constitutively expressed, cytoplasmic protein Fba1. Diploids were sporulated and tetrads dissected to obtain *hsu1*Δ haploids expressing either eGFP (WY2608) or mOrange (WY2612). Colonies expressing the fluorophores show a distinct colour and can therefore be distinguished by visual inspection. Since *hsu1* deletion resulted in the insertion of a KanMX cassette, we removed the resistance gene in some strains to ensure that any fitness difference we observe does not result from the extra KanMX expression in *hsu1*Δ. To do this, we generated a plasmid carrying the dominant selection marker ClonNAT and the *CRE* gene (WSB194). The KanMX cassette which replaced *HSU1* included LoxP flanks and could therefore be looped out by Cre activity. WY2608 was transformed with WSB194 and a selected transformant was allowed to grow to saturation in rich medium (YPD). The culture was appropriately diluted to yield approximately 300 colonies on a YPD plate. Colonies were then simultaneously patched onto YPD plates with and without G418. A colony that could grow on YPD but not on the G418 plate (i.e. had successfully looped out the KanMX cassette) was further propagated by streaking on YPD. A few colonies were selected and simultaneously patched on YPD plates with and without ClonNAT to select for loss of the Cre plasmid. A colony that grew on YPD but not ClonNAT was selected and stored to serve as an eGFP-labelled *hsu1*Δ strain without KanMX (WY2637).

For constructing the strain with *HSU1* tagged at the C-terminus with GFP, the eGFP-KanMX cassette was amplified from WSB65 [63] using primers WSO710 and WSO706. Wildtype S288C (WY2601) and RM11 (WY1203) were transformed with the cassette and transformants were selected on a G418 plate. Transformants were confirmed using a checking PCR with primers WSO711 and WSO159 resulting in strains WY2616 (S288C) and WY2618 (RM11). These strains were mated with MATα *met17*Δ strains (WY577 and WY2533 for S288C and RM11, respectively), and diploids were sporulated. The dissected tetrads were screened on a minimal medium plate as well as a G418 plate to select *met17*Δ spores with *HSU1-eGFP* for each strain background (WY2620 and WY2623).

### Measuring growth dynamics and growth rate

All experiments were initiated with exponentially growing cells for better reproducibility. The evening before an experiment, saturated YPD overnights were used to inoculate minimal medium supplemented with any necessary amino acids for each strain. The next day, optical density at 600 nm (OD) was tracked to check that cells were growing exponentially. Only cultures between 0.2 and 0.4 readings were used to initiate experiments.

For density-dependence assays with *met17*Δ cells, exponentially growing cells were washed thrice in minimal medium before resuspending and appropriately diluting in minimal media to achieve the desired cell densities. For each starting cell density, a single cell suspension was prepared, and equal volumes of this suspension was distributed into sterilized, factory-clean glass tubes.

OD measurements were carried out on a Genesys 20 spectrophotometer (Thermo Fisher Scientific, Waltham, MA, USA) with an adjustable adapter for tubes of different diameters. Each tube was measured thrice with small rotations and the middle value was noted, to avoid the influence of scratches on the glass surface. Any values under 0.001 were below the sensitivity of the machine and were thus replaced by 0.001 for plotting. Note that on this device with tubes of 13-mm diameter, we have estimated that 1 OD corresponds to roughly 7 × 10^7^ cells. On this device, OD no longer scales linearly with cell density beyond 0.5. The data plotted here has not been corrected for this nonlinearity.

For plate experiments, cells were cultured in a Costar 3370 96-well plate at a volume of 150 μl per well. The plates were maintained in a 30ºC incubator and OD was periodically measured with the lid on using a Biotek Synergy MX plate reader. Prior to any measurement, the cultures were agitated to suspend cells using a Teleshake magnetic shaking device (Thermo Fisher Scientific, Waltham, MA, USA) and a custom-built “lid warmer” was used to remove condensation from the plate lid [64]. In each plate, the bottom row was filled with sterile medium for blanking. Any values under 0.001 were below the sensitivity of the machine and were thus replaced by 0.001 for plotting.

For figures S10, S12B and S13, growth rates were measured as the slope of natural log of OD measurements over time. Due to the non-linearity above 0.5 on our spectrophotometer, growth rates were only measured from OD values between 0.2 and 0.5. In S11 Fig, flow cytometry or colony counting was used to measure population densities in co-cultures. In this case the difference in growth rates of the two population was calculated as the slope of natural log of population ratios over time.

### Fluorescence microscopy

The microscopy set-up comprises a Nikon Eclipse TE-2000U inverted fluorescence microscope (Nikon, Tokyo, Japan) connected to a cooled CCD camera for fluorescence and transmitted light imaging. Image acquisition is done with an in-house LabVIEW program. For imaging at different timepoints, 4 μl of the cell culture was spread under a coverslip on a slide. Images were captured using either a 100x objective (Fig 4) or a 40x objective (Fig 5B and S14A Fig). For imaging GFP, the filter cube used was Chroma 49002-ET-EGFP (exciter: 470/40x, emitter: 525/50m, Dichroic: T495LP; Chroma Technology Corp, Bellows Falls, VT, USA). Imaging conditions and exposure times were kept constant for imaging different treatments.

### Flow cytometry

A detailed description has been previously published [65]. Population compositions were measured by flow cytometry using a DxP10 (Cytek, Fremont, CA, USA). To measure absolute cell densities in a sample, fluorescent beads (3-μm red fluorescent beads Cat R0300, Thermo Fisher Scientific, Waltham, MA) of known concentration were added to each sample. The bead concentration was determined by counting on a haemocytometer. Additionally, a final 20,000-fold dilution of the “death dye” ToPro3 (Molecular Probes T-3605; Eugene, OR, USA) was added to each sample to distinguish between dead and live cells [65]. ToPro3-stained dead cells form a distinct cluster from live cells by producing a high signal on the RedFL1 channel (EX: 640nm, filter: 661/16BP). Gating and subsequent analysis were done using FlowJo™ v10.8 Software (BD Life Sciences, Ashland, OR, USA). In brief, beads were first separated from cells based on their high emission on ViolFL1 (EX: 405nm, filter: 445/50BP) and BluFL2 (EX: 488nm, filter: 585/45BP) channels. The cell cluster was further refined by their scattering profile and dead cells were excluded according to the RedFL1 signal. Within live cells, GFP-positive cluster was defined by their BluFL1 (EX: 488 nm, filter: 530/30BP) emission and mOrange-positive cluster was defined by their YelFL1 (EX: 561 nm, filter: 590/20BP) emission.

### Competition

For competition assays, *hsu1*Δ and wildtype strains which constitutively expressed Fba1 tagged with a distinct fluorophore were generated. Each strain was grown to exponential phase in minimal medium and growth rate was tracked for 1-2 hours. The cells were then washed 3 times with sulfur starvation medium, OD values were measured, and the cells were diluted in the starvation medium to yield cultures of 0.1 OD each. 1.5 ml from each culture were combined in factory-clean 13-mm glass tubes to yield a 3ml volume with 1:1 ratio of both genotypes. Three replicate co-cultures were set up within each experiment. The co-cultures were assessed at different timepoints, usually 3-4 times on day one and once a day for subsequent days. For experiments in S11 Fig, samples were collected for both flow cytometry assessment and colony counting at each time point. Flow cytometry samples were prepared as explained in section “Flow Cytometer” and appropriately diluted to yield an event rate of under 1000/s to errors in counting. To achieve consistency in manual colony counting, samples were diluted to get roughly 300 colonies on each plate. Cell densities were however estimated by OD for plating and were therefore not very accurate. In practice, we obtained 150-200 colonies on each plate. Cells were spread on two YPD plates for each time point and allowed to grow for 2-3 days in a 30ºC incubator before counting. A Nikon AZ100 upright microscope was used to distinguish colonies expressing GFP from those expressing mOrange.

As a control, the effect of the fluorophore was tested by swapping the fluorophores on the genotypes and repeating a flow cytometry-based competition assay over the first 8 hours in sulfur starvation medium (S11B Fig). The choice of fluorophores made a measurable impact on the assay (S11C) and we therefore did not use competition assays to determine the fitness difference under high sulfide exposure. Instead, we assessed the impact of sulfide in monocultures of each genotype and assessed cell densities with flow cytometery and colony counting.

### Mathematical model

The basic model and parameter values are described in the main text. The detailed development of the model is provided in Supplementary Text 1, along with the inference of some parameters (*r* and *k*_*s*_) from data-fitting. Sulfide consumption rate *c* was experimentally determined by measuring the increase in cell density at different sulfide concentrations (S15 Fig). Maximum growth rate *g*_*max*_ of *met17*Δ was measured from the 50-μM and 100-μM datasets in the sodium hydroxide experiment (Fig 2F), since growth was the fastest and deterministic under these sulfide treatments. Growth rate was calculated as the slope of natural log of optical density measurements over time. Carrying capacity *K* was also determined from the NaHS experiment by correcting stationary-phase optical density measurements for the non-linearity of our instrument at high OD values and converting the values to cell densities using the relationship 1 OD = 7 × 10^7^ cells. Note that data used for parameter inference and measurement correspond to RM11 yeast.

## ACKNOWLEDGEMENTS

We would like to thank Mark Roth for immensely useful discussions about the chemistry and handling of sulfide, as well providing us with a stock solution of sulfide. We also appreciate discussions with Maitreya Dunham and Dana Miller. We acknowledge the assistance of Olly Jarvis for competition assays, and that of Aurelia Zhu and Michael Gong for generating yeast strains. We are grateful to the Ralser lab for sharing yeast strains, and to Hanadi Rammu for preparing a sulfide stock for us in an anaerobic chamber. Li Xie, David Skelding and all members of the Shou lab have been vital in providing critical input to this work. We thank Shawna Miles for yeast protocols, and Linda Breeden and members of the Sue Biggins lab for hosting and supporting us during our transition from Seattle to London.

