## Supplementary figures for "When is an auxotroph not an auxotroph: how budding yeast lacking *MET17* collectively overcome their metabolic defect"

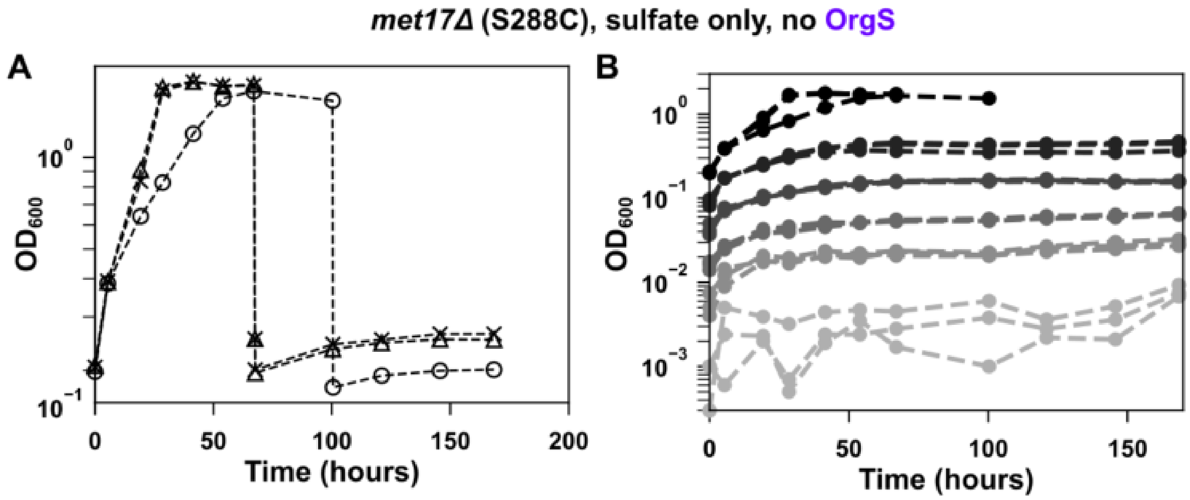


**Supplementary Figure 1: *met17Δ* in the S288C background can grow on sulfate, but growth is less prominent than in the RM11 background**. *met17Δ* (WY576, S288C) cells were grown inliquid SD minimal medium containing sulfate but no organosulfurs. Growth was assessed by measuring optical density at 600 nm ($OD_{600}$) **A**) *met17Δ* can grow to saturation. Each line represents a technical replicate. Unlike in RM11, the saturated cultures do not regrow upon dilution. **B**) *met17Δ* growth on sulfate is density-dependent, but requires a high initial density to achieve saturation. Darker gray shades indicate higher initial cell densities, and three technical replicates were started at each density. Only cultures at the highest initial cell density tested ($OD_{600}$ = 0.2) grew to saturation. Cultures in A are the same as the highest density cultures in B. Cultures were of 7 ml volume in glass tubes of 18-mm diameter with loosely fitted plastic lids.

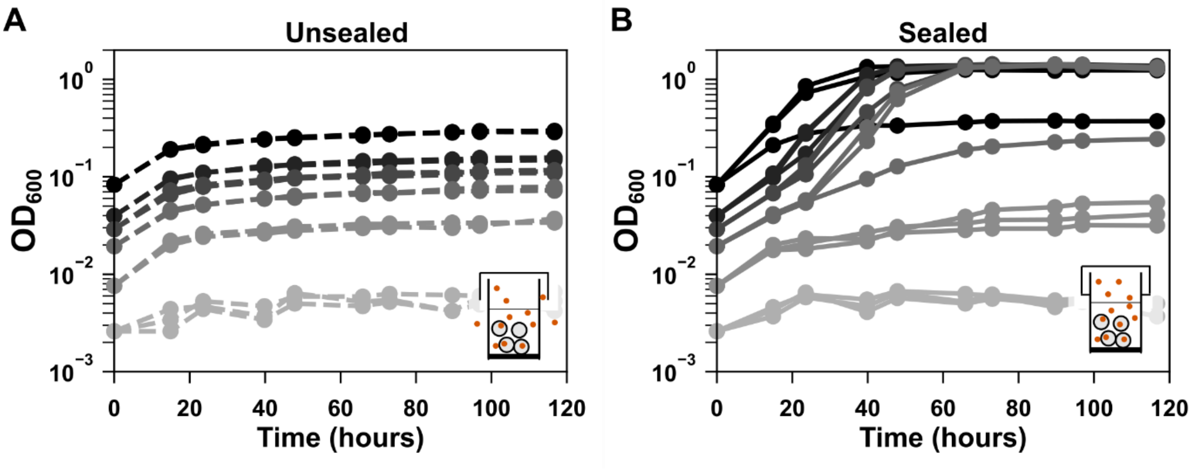


**Supplementary Figure 2: Lowering gas escape increases the propensity of S288C *met17*∆ to grow on sulfate**. *met17*∆ (WY576) cultures were initiated in minimal medium SD at different initial cell densities (denoted by different shades of gray). At each density, 6 replicate cultures of 2.5 ml each were set up in glass tubes with plastic lids. Of these, 3 tubes were additionally sealed with parafilm to lower gas escape, while the remaining 3 only had plastic lids that allow gas exchange. **A**) Unsealed tubes showed only residual growth at all densities, unable to reach saturation. **B**) In contrast, in sealed tubes, growth was observed at initial cell densities of $OD_{600}$ 0.02 and above, and stochasticity was evident as the three replicates at a given initial cell density showed divergent growth dynamics.

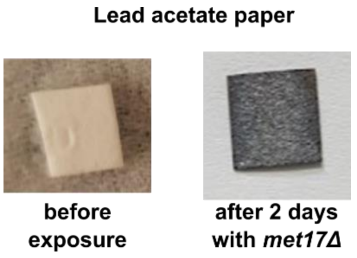


**Supplementary Figure 3: Hydrogen sulfide gas is released by *met17*Δ growing on sulfate.** A square of lead acetate paper placed in between the wells of a 96-well plate where *met17*Δ were growing in minimal medium turned black after 2 days. Black colour indicates the formation of lead sulfide upon reaction of lead acetate with hydrogen sulfide. Strain: WY2531.

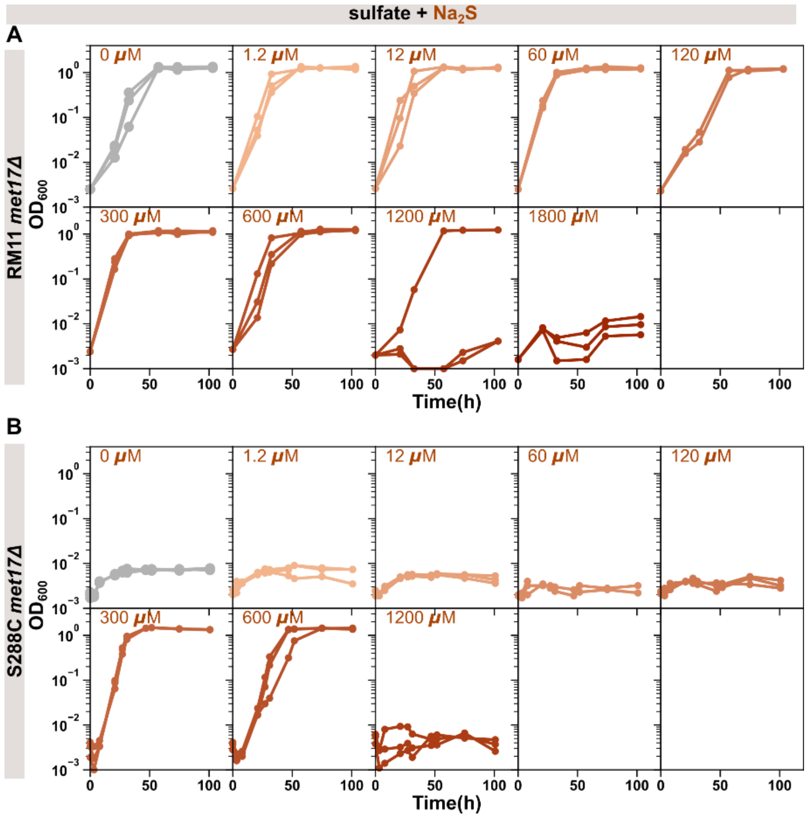


**Supplementary Figure 4: Sodium sulfide promotes the growth of *met17*Δ in sulfate over a range of concentrations, but becomes toxic at high concentrations.** Different concentrations of sodium sulfide ($Na_2S$) were added to low-density cultures of *met17*Δ in minimal medium. Three technical replicates were set up for each sulfide concentration and tubes were additionally sealed with parafilm and cling film to prevent the loss of sulfide. (**A**) Growth curves of RM11 *met17*Δ (WY2548). Note that even without additional sulfide, the cultures grew to saturation because the additional sealing reduced escape of sulfide released by the cells (top left in A, gray lines). Addition of up to 600 µM of $Na_2S$ sped up cell growth, i.e growth curves reached saturation at an earlier timepoint than in the absence of additional $Na_2S$ (grey). The curves at 120 µM are an exception that show slower growth but this was not a consistent observation. Concentrations of 1.2 mM and above could inhibit growth. (**B**) Growth curves of S288C *met17*Δ (WY2590). Growth promotion of *met17*Δ on sulfate is only observed with addition of 300 or 600 µM $Na_2S$. The highest concentration tested (1.2 mM $Na_2S$) impaired growth.

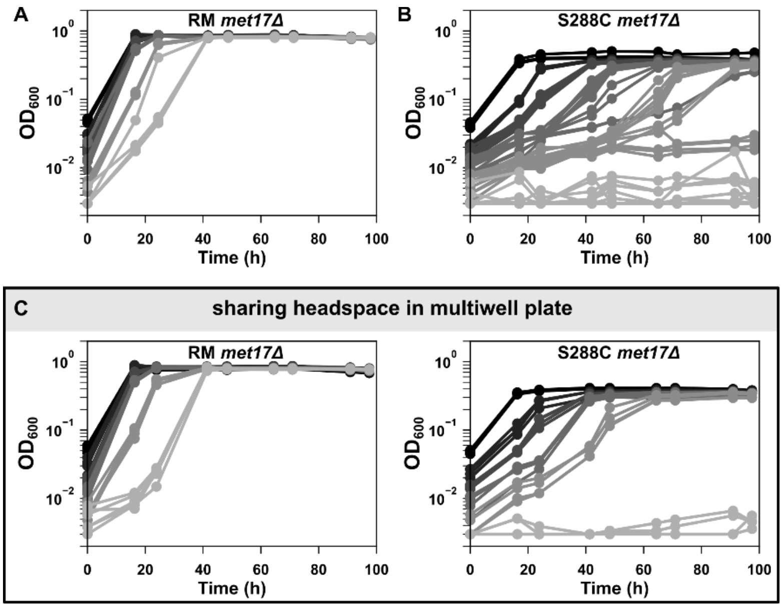


**Supplementary Figure 5: S288C *met17*Δ release lower $H_2S$ than RM11 *met17*Δ.** Different initial cell densities of *met17*Δ were grown in liquid minimal medium in separate wells of a 96-well plate, thus sharing headspace. (**A**) All densities of RM11 *met17*Δ (WY2531) grew to saturation within 40 hours. (**B**) In contrast, S288C (BY) *met17*Δ (WY2515) required much longer times to grow to saturation, and lower cell densities never grew to saturation. (**C**) When the two strains occupied different wells of the same multi-well plate such that they shared headspace and volatiles, all (except the lowest cell density) of S288C *met17*Δ grew faster (compare with B). Since growth propensity increases with increasing sulfide concentrations, the growth promotion of S288C in the vicinity of RM11 indicates that RM11 *met17*Δ release more H2S than S288C *met17*Δ. For the S288C strain, SD medium was additionally supplemented with histidine, leucine, and uracil to cover the nutritional requirements caused by additional (engineered) auxotrophic mutations.

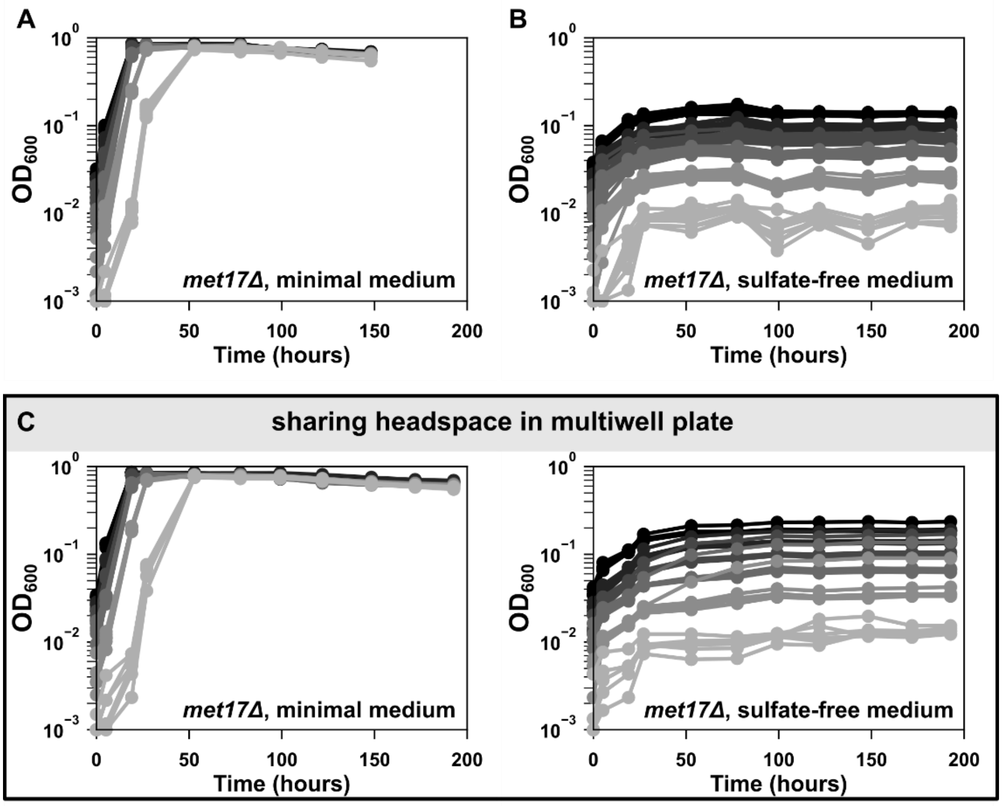


**Supplementary Figure 6: Growth of *met17*Δ in minimal medium requires inorganic sulfate.** While all densities of *met17*Δ (WY2548, RM11) could grow to saturation in minimal medium containing sulfate when sharing headspace in multiwell plates (**A**), *met17*Δ could not grow in sulfate-free medium at any cell density (**B**; see Methods). Even when these two sets of populations shared headspace in the same multiwell plate, *met17*Δ could not grow in sulfate-free medium, suggesting that sulfide from neighbouring wells is insufficient to support the growth of *met17*Δ and they require the sulfide they themselves produce by sulfate reduction. This is in contrast to *met14*Δ that are unable to produce their own sulfide, but could grow on sulfide from neighbouring wells (Fig 2E). One possible explanation is that while *met17*Δ can grow by assimilating sulfide, their mechanism of utilizing sulfide is of low efficiency compared to the Met17-dependent sulfide assimilation observed in *met14*Δ*.*

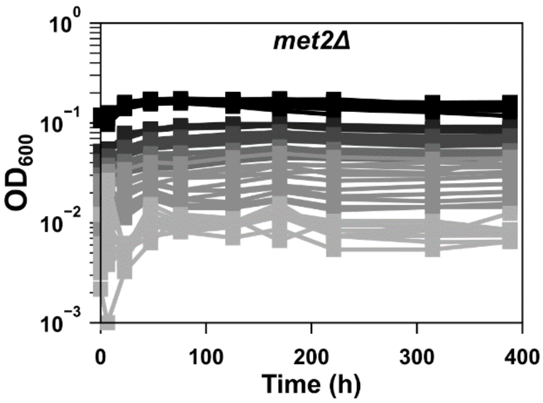


**Supplementary Figure 7: *S. cerevisiae* cannot bypass the need for *MET2* to assimilate inorganic sulfate.** *met2*Δ (WY2538, RM11) could not grow in minimal medium at any initial cell density. Higher initial cell densities are denoted by darker shades of gray. All populations were sharing headspace in a multiwell plate to favour the exchange of $H_2S$ and maximize the chances of growth by a sulfide-dependent alternative mechanism.

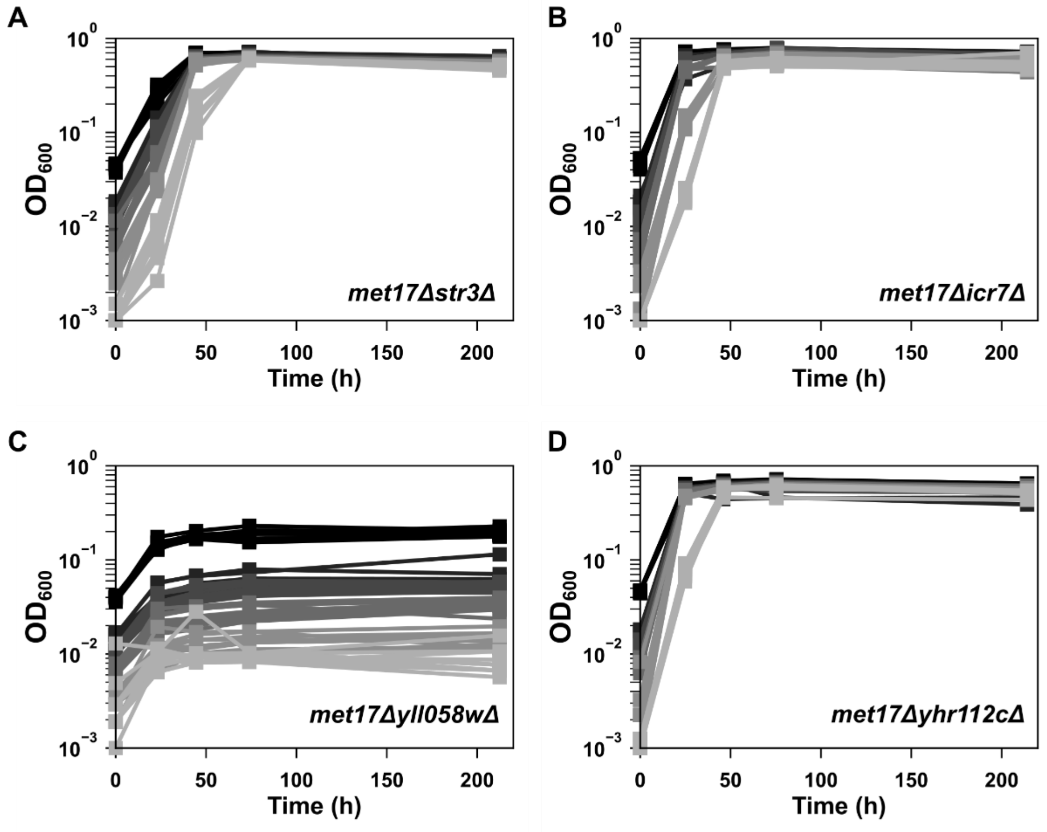


**Supplementary Figure 8: Identifying the gene responsible for assimilating sulfide in *met17*Δ.** Four candidates, selected on the basis of sequence similarity to Met17, were tested through the double mutant screen described in Fig 3B. In brief, we were looking for the gene which when deleted along with *MET17* completely abrogated growth on sulfate (i.e. in minimal medium without organosulfur supplements). Strains belonged to the BY4741 (S288C) Yeast Deletion Library which, along with a *met17* deletion, also lack synthesis of histidine, leucine and uracil. Minimal medium was thus supplemented with these three nutrients, but methionine was removed once the cells grew to exponential phase. Cultures were diluted to different cell densities (denoted by different shades of gray), and grown in a multiwell plate to maximize the chances of sulfide-dependent population growth of *met17*Δ. Of the candidates tested, only *YLL058W* passed the screen with only residual growth observed at all cell densities (C). Strains: WY2584 (*str3*Δ), WY2587 (*icr7*Δ), WY2586 (*yll058w*Δ) and WY2585 (*yhr112c*Δ).

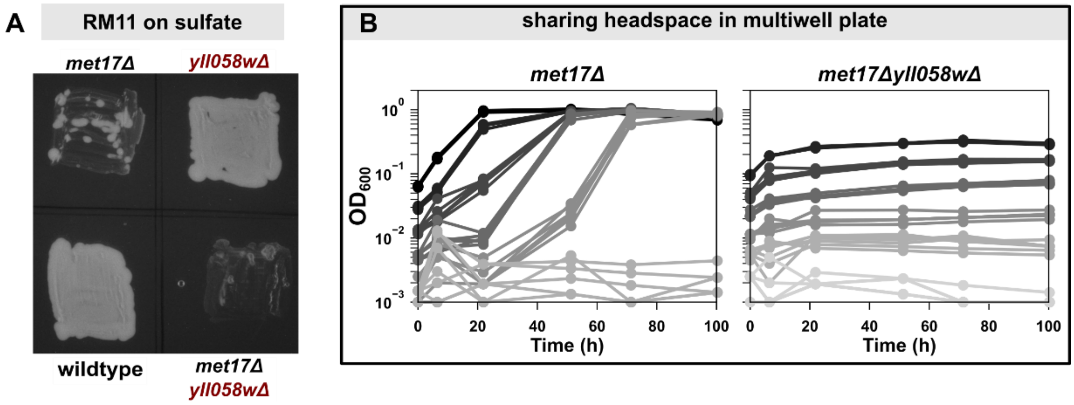


**Supplementary Figure 9: Sulfide assimilation is defective in *met17Δyll058wΔ* double knockouts. A**) The double mutants *met17Δyll058wΔ* in RM11 background (WY2642) do not show papillae when patched onto agar plates containing SD minimal medium, even while growing on the same plate as $H_2S$-releasing *met17Δ* (WY2548). *yll058wΔ* (WY2639) grow dense lawns similar to wildtype RM11 (WY2641). All four genotypes were patched onto the same agar plate which was sealed with parafilm and imaged after 3 days. **B**) Double mutants (WY2595, S288C) were inoculated in minimal medium at different initial cell densities in the same multi-well plate where different densities of *met17Δ* (WY2590, S288C) were growing. This should allow *met17Δyll058wΔ* to consume $H_2S$ released by *met17Δ*, as was observed for *met14Δ* in Fig 2E. However, the double mutant could not grow indicating that they are impaired in sulfide assimilation even when sulfide was supplied.

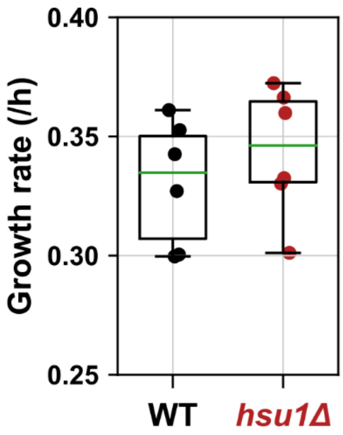


**Supplementary Figure 10: Exponential growth rates of *wildtype* and *hsu1Δ* are comparable.** Growth rates of S288C wildtype (WT; WY1376, WY1377, WY2601) and *hsu1Δ* yeast (WY2597, WY2608, WY2637) during exponential growth in minimal medium are not significantly different (p-value = 0.19, two-tailed paired Student's t-test, where growth rates measured on the same day were paired). Growth rates were calculated as the maximal slope of natural log of optical density measurements against time. Box plot shows quartiles, with whiskers extending to the most extreme datapoints and median marked as the green line.

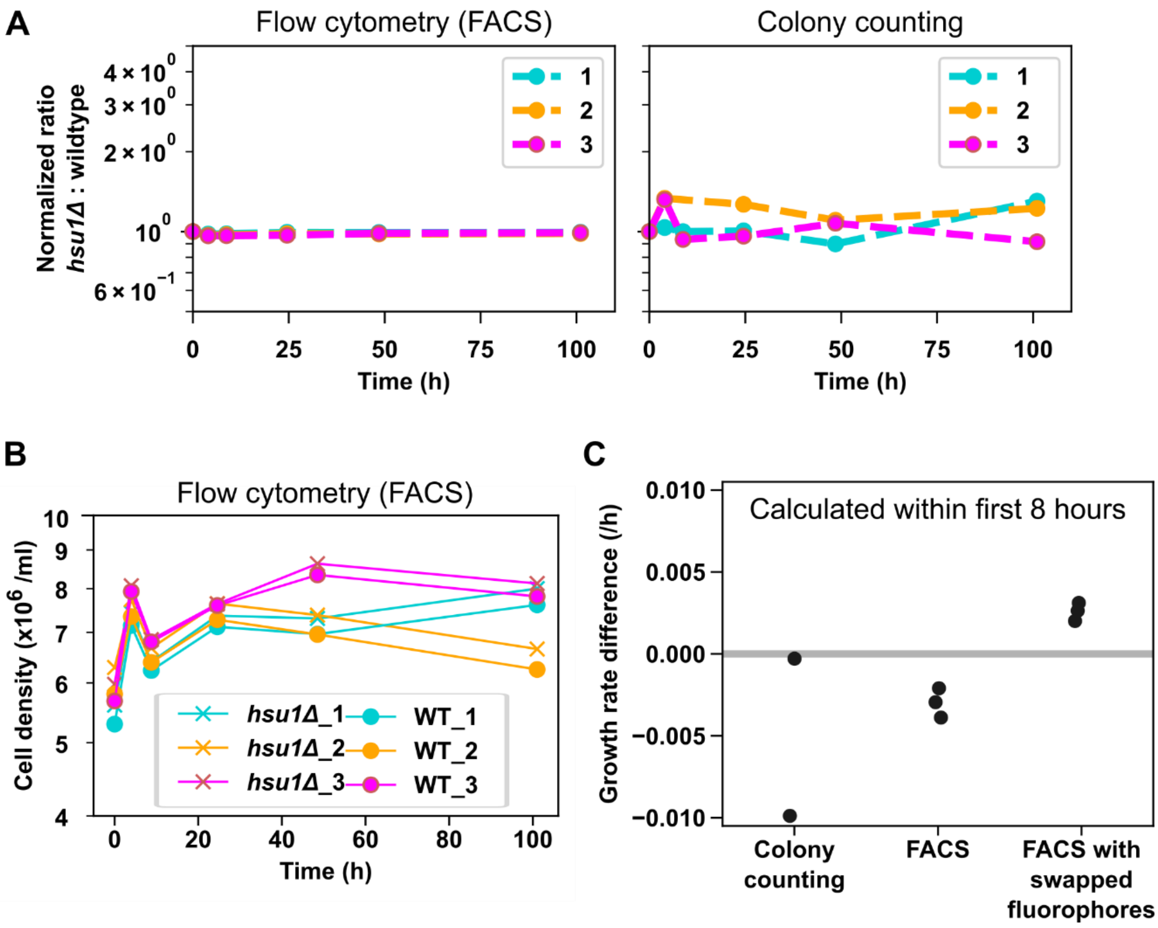


**Supplementary Figure 11: No disadvantage detectable for *hsu1*Δ under sulfur starvation. A, B**) Competition assays were performed with wildtype (WY1377) and *hsu1*Δ (WY2637) yeast under sulfur starvation by mixing the two genotypes at a 1:1 ratio in sulfate-free medium and following population dynamics using either flow cytometry or colony counting. Each genotype carried a distinct fluorescent protein to allow identifying the population. Three replicate mixtures were set up in each experiment and sampling was done for both flow cytometry and colony counting at each timepoint. Samples were appropriately diluted to get single events on flow cytometry and for manually counting colony forming units on a rich medium plate. Ratios were normalized to initial value. The ratio remained largely stable in flow cytometry data (A, left) whereas large variability with no distinct trend was observed in colony counting (A, right). Colony counting is inherently noisy as counting can only be done on the order of hundreds, whereas flow cytometry measures tens of thousands of events at each time point. Population dynamics measured by flow cytometry (B) shows that although there were variations among tubes, within a tube the population sizes of two genotypes were very similar. The initial residual growth was enabled by cellular sulfur store. **C**) Growth rate difference between *hsu1*Δ and wildtype was measured as the slope of natural log of ratios in co-culture experiments over the first 8 hours. For each condition, each dot is a replicate from one competition experiment. Swapping fluorophores on the genotypes led to a small positive deviation, whereas the original pairing produced a small negative deviation, suggesting a minor contribution from such experimental details. Overall, the fitness difference between wildtype and *hsu1*Δ, if any, is too small to be detected in our assay. As expected, colony counting is far noisier than flow cytometry. Colony counting and flow cytometry data points in C were from the same dataset depicted in A and B. Strains for swapped fluorophores were WY2612 (*hsu1*Δ) and WY1364 (wildtype). All strains are of S288C background.

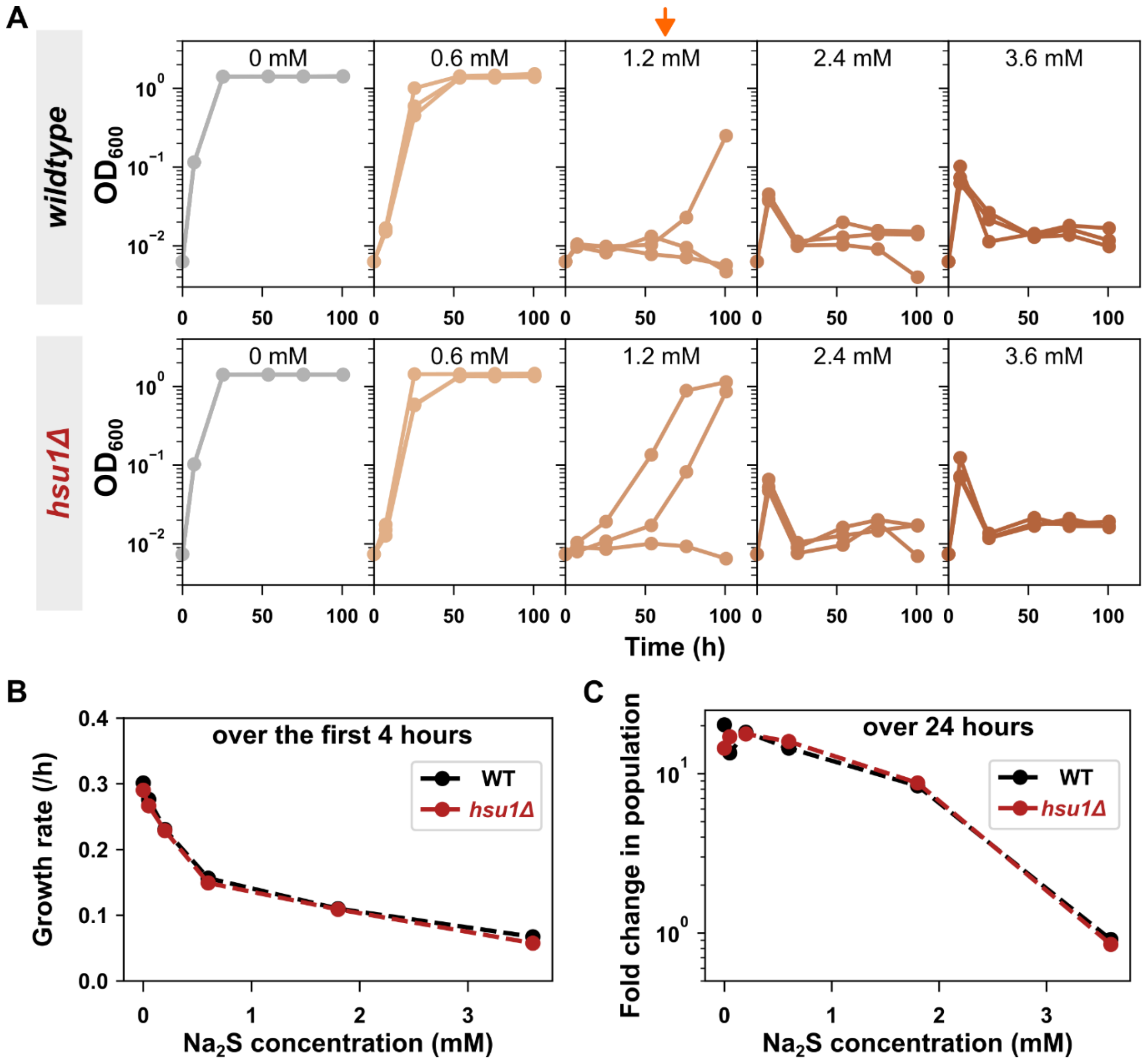


**Supplementary Figure 12: *hsu1*Δ and wildtype behave similarly to high sulfide exposure. A**) The concentration where sulfide impairs growth is similar for *hsu1*Δ (WY2597) and wildtype (WY2601) populations. The effects of different concentrations of $Na_2S$ on yeast growing in SD minimal medium were compared by measuring population dynamics using optical density. For both genotypes, at 1.2 mM sulfide, at least one of three replicate populations failed to grow (highlighted with orange arrow). **B**) Sulfide impacts the growth rate of *hsu1*Δ and wildtype in a similar way. Growth rate was measured as the slope of natural log of OD measurements over the first four hours of sulfide exposure in 2.5-ml minimal medium cultures. Increasing concentrations of $Na_2S$ led to a decrease in growth rate, but the response in both genotypes was identical. **C**) Both *hsu1*Δ and wildtype populations had similar population dynamics during 24 hours of sulfide exposure. Population size was measured using flow cytometry before adding $Na_2S$ and after 1 day of growth in different concentrations of $Na_2S$. Datasets **B** and **C** come from the same experiment with strains WY2637 (*hsu1*Δ) and WY1364 (wildtype). All strains are of S288C background.

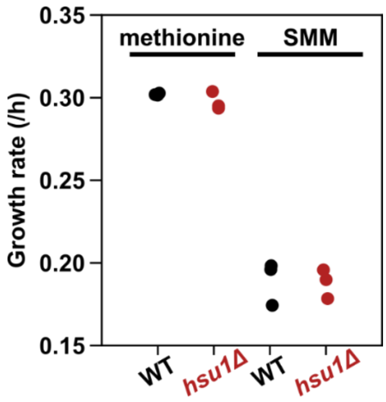


**Supplementary Figure 23: *hsu1*Δ and wildtype show a similar growth rate on S-methylmethionine.** The growth rates of *hsu1*Δ (WY2597) and wildtype (WY2601) were indistinguishable in liquid sulfate-free medium supplemented with either methionine or S-methylmethionine (SMM). Both genotypes had a slower growth rate on SMM as compared to methionine. Growth rates were calculated as the slope of natural log of optical density measurements over the first 8 hours of growth on the organosulfur. Each dot represents a biological replicate, i.e. a different colony from the same yeast strain. Both strains are of S288C background.

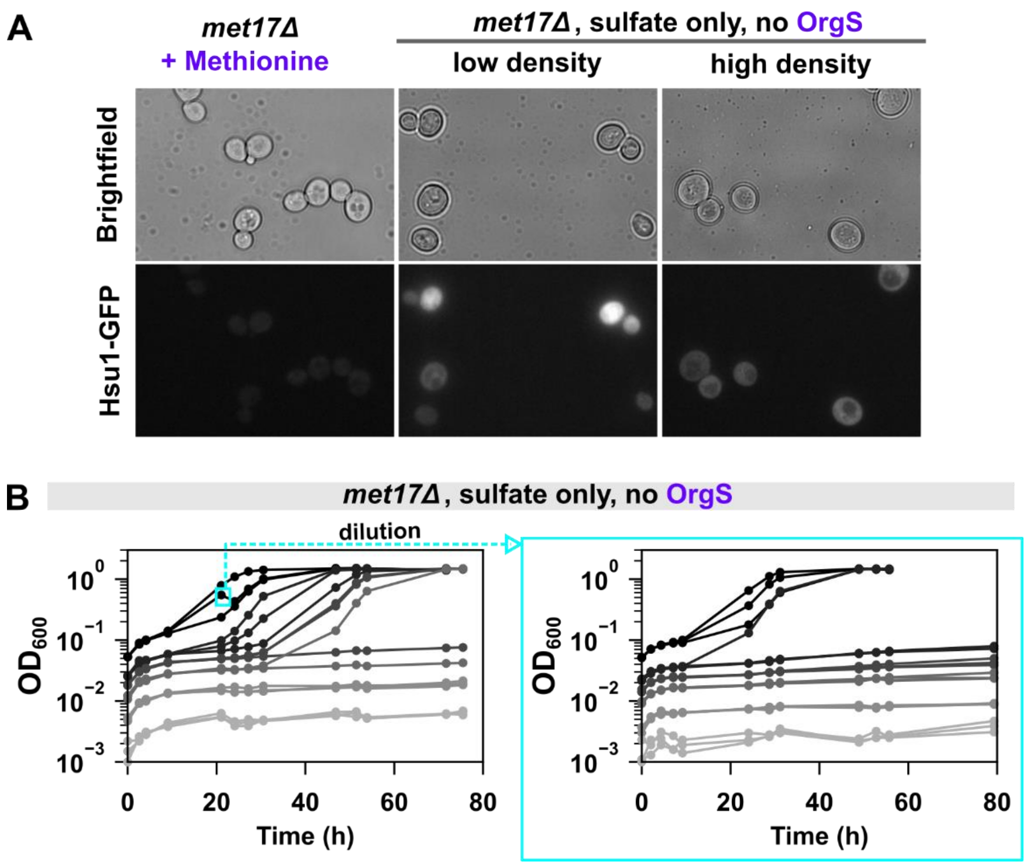


**Supplementary Figure 14: A cell-state switch does not underpin the stochastic growth dynamics of S288C *met17*Δ in sulfate. A**) The expression of *HSU1-GFP* in S288C *met17*Δ (WY2620) does not correlate with growth status. While exponential cells growing in methionine do not show considerable expression of Hsu1-GFP (left panels), cultures inoculated at both low and high cell densities show expression within 4 hours of transfer to SD minimal medium containing sulfate, but no organosulfurs. These cultures did not eventually grow to saturation, indicating that if a cell-state switch results in the stochastic growth dynamics, the expression of Hsu1-GFP cannot be the mechanism of the switch. **B**) If a cell-state switch exists, *met17*Δ that have started to grow on sulfate (cyan rectangle in the left panel) should already have switched on. If density-dependence relies on the cell-state switch, such cells should grow even at low-cell densitiest when re-inoculated into SD medium (right panel). However, this was not observed, suggesting that density-dependence does not result from a cell-state switch or that the switch is too transient to persist through the dilution. Strain: WY2590.

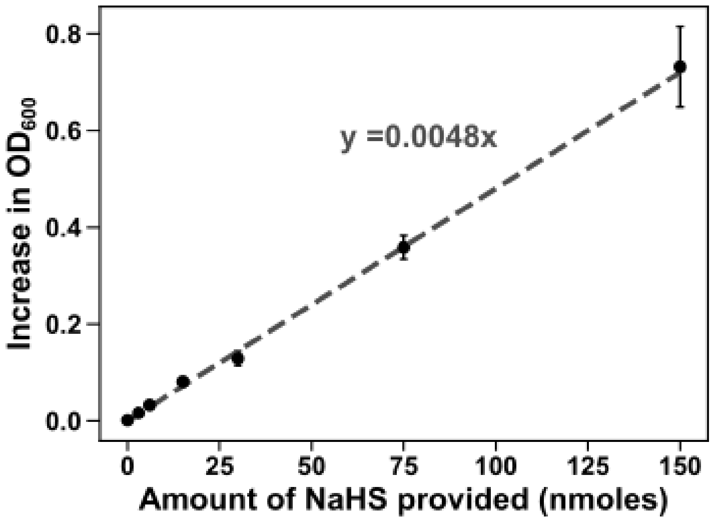


**Supplementary Figure 15: Consumption rate of sulfide $c$ is approximately 3 fmole/cell birth in RM11 yeast.** RM11 *met14*Δ yeast (WY2539) were grown to exponential phase in minimal medium supplemented with methionine and washed and transferred to minimal medium without organosulfurs. This genotype was chosen as it could not consume the sulfate in minimal medium (Fig 2D). Cells were starved for 24 hours before addition of different amounts of sodium hydrosulfide (NaHS) to three replicate cultures each. An end-point optical density measurement was done after 3 days and the increase in $OD_{600}$ was plotted against the amount of NaHS provided. The slope of the linear regression (0.0048) was used to calculate the amount of sulfide consumed per cell birth. 1 unit of $OD_{600}$ corresponds to $7 \times 10^7$ cells in our set-up.

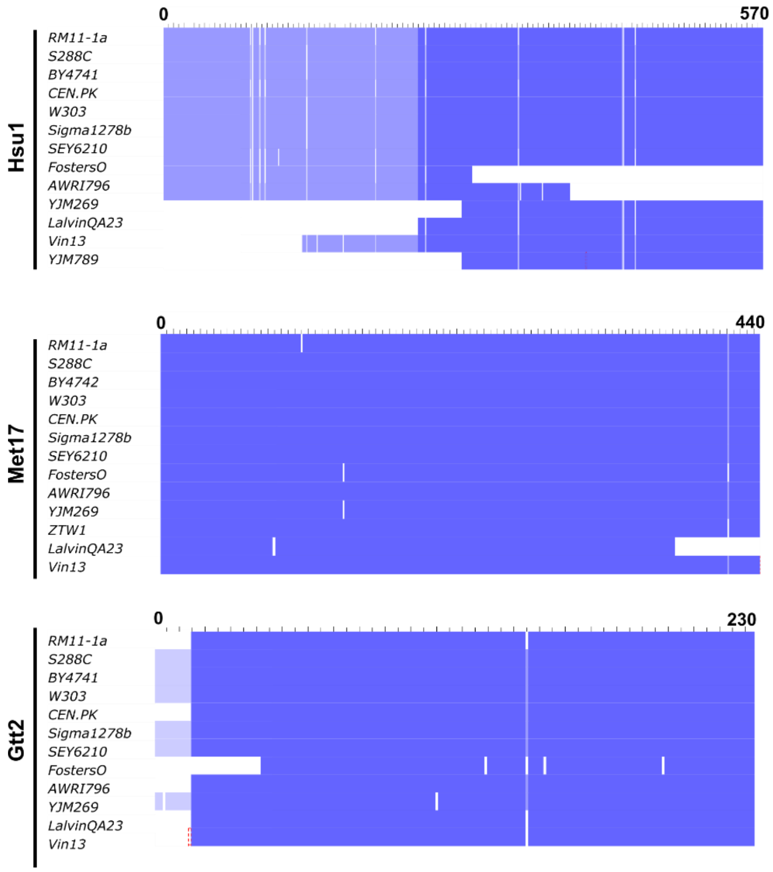


**Supplementary Figure 16: Hsu1 proteins shows a high degree of polymorphism among yeast strains.** Protein sequences from 41 strains in the *S. cerevisiae* Genome Database were aligned for the protein Hsu1, Met17 and Gtt2. Higher percentage identity is denoted by darker blue shades. A selection of the strains are shown here to highlight the high degree of polymorphisms tolerated in Hsu1, compared to Met17. Notably, large deletions were observed in Hsu1 sequences from some strains. HSU1 is located close to the telomere on Chromosome XII. However, GTT2 which is more proximal to the telomere shows fewer regions of variability. Variations in the initial stretch may result from erroneous annotation of start codon. Altogether, these data suggest that the function of HSU1 may be less important to the cell's fitness than that of Met17 or Gtt2.
