## Supplementary Text for "When is an auxotroph not an auxotroph: how budding yeast lacking *MET17* collectively overcome their metabolic defect"

### Supplementary Text 1: Details of model assumptions and parameter fitting

#### Contents

1 **A dynamic model for cell density and $H_2S$ concentration** 1

2 **Fitting unknown model parameters ($k$ and $r$)** 1

3 **Our model describes the density-dependent behavior of *met17Δ*** 4

4 **Timescale separation avoids the need to explicitly model gas phase** 5

### 1 A dynamic model for cell density and $H_2S$ concentration

We used a model in which cells both produce $H_2S$ at a fixed rate, and also consume $H_2S$ as a biological building material:

$$\frac{dx}{dt} = \frac{g_{max}s}{k+s}\left(1 - \frac{x}{k_{cap}}\right)x$$
$$\frac{ds}{dt} = rx - \frac{dx}{dt}c - \delta s$$

The state variables $x$ and $s$ are cellular density and $H_2S$ concentration respectively at time $t$. The parameters are defined below:

- $g_{max}$ is the maximum growth rate.
- $k_{cap}$ is the cellular carrying capacity.
- $k$ is the "Monod constant", i.e. the concentration of $H_2S$ at which growth rate is half-maximal.
- $r$ is the rate at which a cell releases $H_2S$.
- $c$ is the amount of $H_2S$ consumed to produce a new cell.
- $\delta$ is the rate at which $H_2S$ is lost from the culture tube due to gas leakage.

One possible concern that comes to mind is that this model does not explicitly include the phase transition of hydrogen sulfide between the aqueous and gas phases. A more complicated model would maintain three state variables, one for cell density, and two for the two different states of $H_2S$ (aqueous versus gas) . However, in section 4, we show that our simpler model is actually sufficient to capture cell-density dynamics (despite the existence of two $H_2S$ phases), as long as the dynamics of phase transitions of $H_2S$occur on a faster timescale than the other processes involving $H_2S$ (i.e. cellular consumption, cellular release, and leakage).

### 2 Fitting unknown model parameters ($k$ and $r$)

In brief, various values of the Monod constant $k$ and the release rate $r$ were selected, and the model was run with these values and compared to data from experiments where RM11 *met17Δ* yeast were subjected to different concentrations of sodium hydrosulfide (NaHS, which releases $H_2S$). The best fit was obtained with $k = 7.1$ µM and $r = 0.39$ fmole/cell/hr. Note that "fmole" is short for "femtomole".

**Data inclusion criteria:** We fit the model to the data presented in Figure 2F. Trials with an initial NaHS concentration of 0, 1, 10, or 50 µM were included. Trials with a greater initial NaHS concentration were excluded because greater concentrations appear to be toxic to cells, and this toxicity is not included in the model. Also excluded were trials wherein optical density did not reach 1 by the end of the experiment. Measurements after 100 hours were also excluded. This is because after 100 hours, cultures saturated and began to decline in population, but cell death was not included in the model.

**Units:** Optical density (OD) values were converted to cell density according to the following experimentally-determined formula to correct for the fact that higher OD values deviate from linearity:

$$x = \begin{cases} 7 \times 10^7 C(\omega) & \omega > 0.5 \\ 7 \times 10^7 \omega & \omega \leq 0.5 \end{cases}$$

$$C(\omega) = 2.282\omega/(2.748 - \omega)$$

$\omega$ is OD, $C(\omega)$ is "corrected OD" for values above 0.5, and $x$ is cell density in cell/ml. These values are specific for our spectrophotometer when used with 13-mm culture tubes.

**Fitting procedure:** The following parameters were used for fitting the model:

- $g_{max} = 0.26/\text{hr}$
- $c = 3\text{fmole/cell}$
- $k_{cap} = 1.6 \times 10^8 \text{cell/ml}$
- $\delta = 0/\text{hr}$

$g_{max}$, $c$ and $k_{cap}$ were experimentally determined (see Methods: Mathematical model), whereas loss was assumed to be negligable as a simplification. We scanned through different values of $k$ and $r$ to see which combination of values minimized the model error. We varied $k$ from 0.1 to 15 µM (in steps of 0.1 µM), and then from 15 to 50 µM (in steps of size 1 µM). We varied $r$ from 0 to 0.7 fmole/cell/hr (in steps of size 0.01 fmole/cell/hr), and then from 0.7 to 20.2 fmole/cell/hr (in steps of size 0.5 fmole/cell/hr). That is,

$$k = 0.1, 0.2, ..., 14.9, 15, 16, ..., 49, 50 \text{ µM}$$

$$r = 0, 0.01, ..., 0.69, 0.7, 1.2, ..., 19.7, 20.2 \text{ fmole/cell/hr}$$

the model was initialized at the first time point (zero hours) with the initial cell density given by the measured cell density at zero hours and the initial dissolved $H_2S$ concentration given by the concentration of NaHS. The model was "run" via numerical integration, and the numerical solution was compared with the observed cell density values. The goodness of fit to the data was quantified as the root mean squared log-error (RMSLE):

$$RMSLE = \sqrt{\frac{1}{nm}\sum_{i=1}^{n}\sum_{j=1}^{m}\left(\log_2(x_{i,j}) - \log_2(\hat{x}_{i,j})\right)^2}$$

where $x_{i,j}$ is the observed cellular density in the $i$th trial at the $j$th measurement time, and $\hat{x}_{i,j}$ is the same, but in the model. $n$ is the number of trials and $m$ is the number of measurements per trial. Base-2 logarithm is used because, as shown below, it captures the error range nicely. RMSLE indicates the average logarithmic deviation between the model-predicted cellular density and the experimentally measured cellular density. For instance, an RMSLE of 1 indicates that on average, the model-estimated cellular density is either twofold above or twofold below the experimentally measured cellular density. Fig ST-1 shows contours of RMSLE across the two-dimensional grid of parameters. To provide some visual intuition for parameter sensitivity, Fig ST-2 shows how closely the model aligns with the data given either the best-fit parameters ($RMSLE = 0.88$) or a suboptimal parameter choice ($RMSLE = 1.54$).

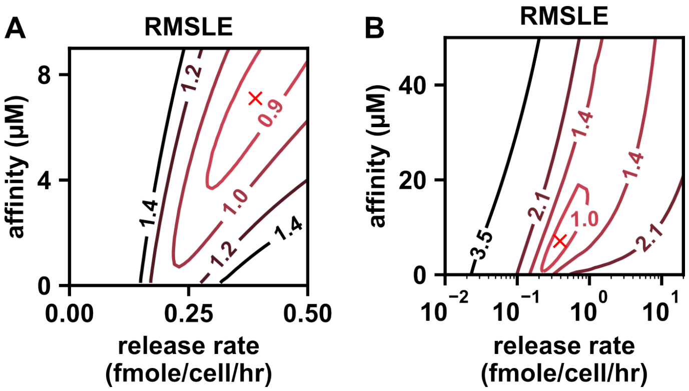


Figure ST-1: Root mean squared log-error (RMSLE) a function of the release rate and Monod constant. (**A**) is zoomed in and (**B**) is zoomed out, with a logarithmic horizontal axis. Contour labels indicate RMSLE. A red "×" marks the position of minimum error, which corresponds to $k = 7.1$ μM and $r = 0.39$ fmole/cell/hr. With these best-fit parameters, the model has an RMSLE of 0.88.

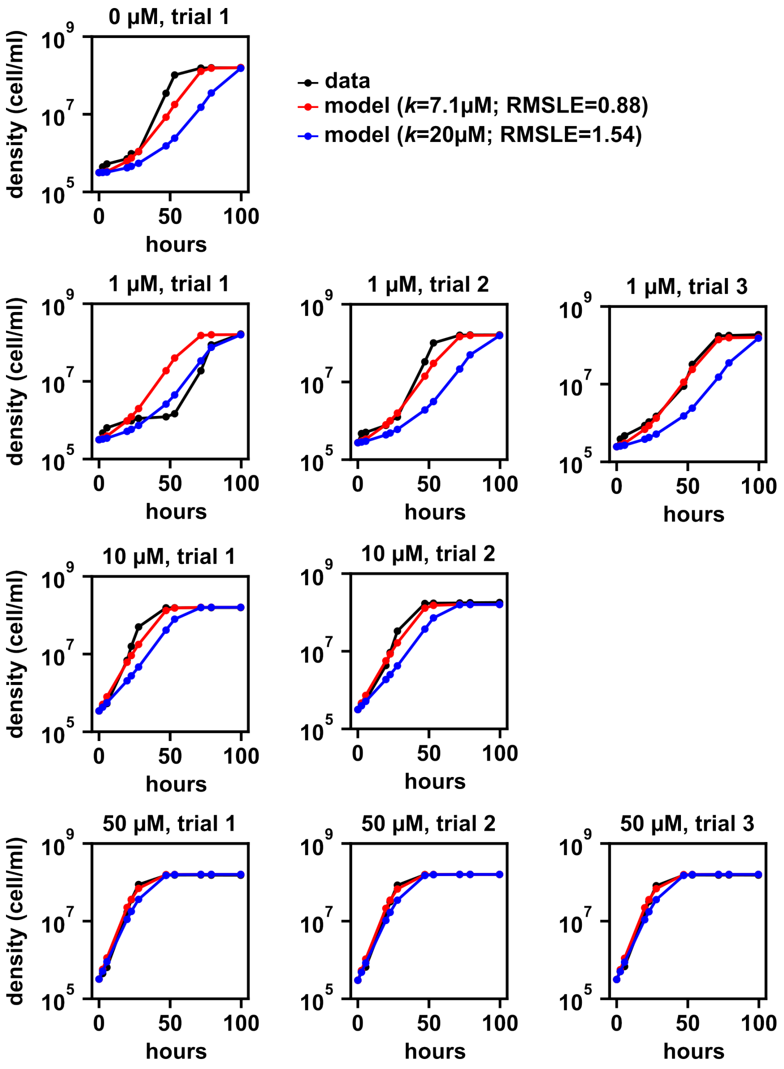


Figure ST-2: Visual comparison of experimentally observed cellular density versus model-estimated cellular density (for two different parameter choices) at various initial NaHS concentrations. Initial NaHS concentration is shown above each chart. Observed densities (i.e. the same data as in Figure 2F in the main text) are shown in black. Red points show model-estimated cellular density with best-fit parameters ($k$ = 7.1 µM and $r$ = 0.39 fmole/cell/hr). Blue points show model-estimated cellular density with $k$ = 20 and best-fit $r$. A comparison between the red and blue curves illustrates how sensitive the fit is to a change in $k$. There is only one chart shown for the initial NaHS concentration of 0 µM because in the other two trials with 0 µM, cells failed to reach an optical density of 1, so these trials were excluded from fitting. One trial with an initial NaHS concentration of 10 µM was excluded for the same reason.

#### 3 Our model describes the density-dependent behavior of *met17*Δ

In the main text, we saw that the growth behavior of *met17*Δ differed from prototrophic yeast in two ways. First, *met17*Δ (but not prototrophs) have density-dependent lags in the growth dynamics. Second, at a given initial cell density, substantial tube-to-tube variation was observed in the lag times for *met17*Δ.

Our model, with the parameters chosen in section 2, provides a possible explanation for the two *met17*Δ phenomena (Fig ST-3A). As in *met17*Δ, there is a longer lag phase at a lower initial density. Moreover, the tube-to-tube variation in the lag phase duration may be due to a difference among culture tubes in the leakage rate $\delta$. Indeed, the model features different lag-phase durations depending on the leakage rate (but only at sufficiently low initial cell density). Additionally, variation in the lag phase does not seem to be attributable to the kind of small fluctuations in initial cell density that might be expected from pipette error (Fig ST-4).

Unlike *met17*$\Delta$, prototrophic strains have no detectable lag at any observed cell density (e.g. *hsu1*$\Delta$ in Fig 3C middle panel). What might explain the difference? One possibility is a difference in the Monod constant. In fact, a ten-fold reduction in the Monod constant can considerably reduce the lags (Fig ST-3B).

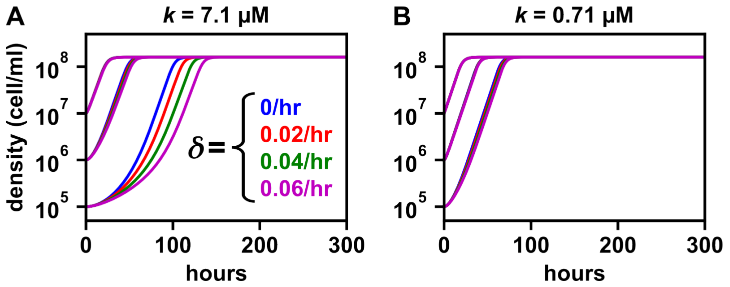


Figure ST-3: The model with fitted parameters displays a lag time that depends on initial cell density and $H_2S$ leakage, and this dependence is reduced or eliminated when different parameters are used. (**A**) With fitted parameters, lag time depends on initial cell density and leakage rate $\delta$. Blue, red, green, and purple denote $\delta$ values of 0, 0.02., 0.04, and 0.06/hr respectively. All curves use the parameter values $c = 3$ fmole/cell; $k_{cap} = 1.6 \times 10^8$ cell/ml; $g_{max} = 0.26/hr$; $r = 0.39$ fmole/cell/hr. The initial $H_2S$ concentration is set to 0 M. (**B**) As in **A**, but where the Monod constant $k$ has been decreased by 10-fold.

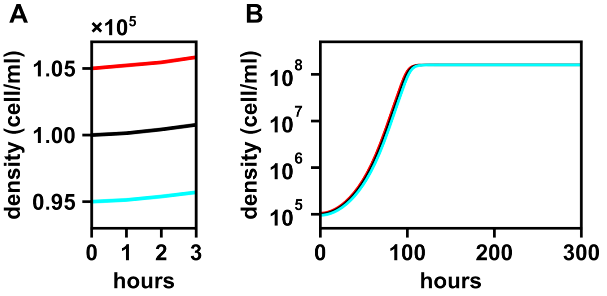


Figure ST-4: Minor variation in initial cell density does not appreciably affect the lag phase according to the model. (**A**) The first three hours of growth according to the model with three different initial densities reflecting a target density of $10^5$ with a possible pipetting error of $\pm 5\%$. The initial $H_2S$ concentration was set to 0 M. The best-fit model parameters were used (i.e. the same model parameters as in Fig ST-3A), and no $H_2S$ leakage was assumed. (**B**) The same three curves, but now shown over the full range of growth. The different initial densities do not lead to appreciably different dynamics, so the curves appear superimposed.

### 4 Timescale separation avoids the need to explicitly model gas phase

In this section, we show that if the kinetics of the aqueous-gas phase transition in $H_2S$ are much faster than the kinetics of other $H_2S$-related processes (i.e. cellular consumption, cellular release, and leakage), then it is not necessary to explicitly include a state variable for gas-phase $H_2S$. To do so, we begin with a "full model" (including both aqueous and gas-phase $H_2S$, and show how it is equivalent to a simpler model (including aqueous $H_2S$ only, but with some parameter adjustments).

For the full model, we assume that Henry's law applies, so that the number of moles of $H_2S$ in aqueous phase is proportional to the number of moles of $H_2S$ in gas phase. Thus if there are $n_{tot}$ moles of $H_2S$ in total, there are $f_{aq}n_{tot}$ moles of $H_2S$ in the aqueous phase and $(1 - f_{aq})n_{tot}$ moles in the gas phase. Furthermore, we use an approximation which says that the aqueous-gas phase transition kinetics are so fast that they appear instantaneous from the point of view of biological activities and $H_2S$ leakage. In this view, if $n_{cons}$ moles of $H_2S$ are consumed in a period of time, we model this by saying that $n_{cons}f_{aq}$ moles are consumed from the aqueous phase, and $n_{cons}(1 - f_{aq})$ moles are consumed from the gas phase in that period of time. A similar approximation applies for $H_2S$ release and leakage. To simplify calculations, we initially let the $H_2S$ state variables denote the number of moles rather than concentration (molarity).

$$\begin{aligned}
\frac{dx}{dt} &= \frac{g_{max}n_{aq}/V_{liq}}{k_{aq} + n_{aq}/V_{liq}}\left(1 - \frac{x}{k_{cap}}\right)x \\
\frac{dn_{aq}}{dt} &= rxV_{liq}f_{aq} - \frac{dx}{dt}cV_{liq}f_{aq} - \delta_{gas}n_{gas}f_{aq} \\
\frac{dn_{gas}}{dt} &= rxV_{liq}(1 - f_{aq}) - \frac{dx}{dt}cV_{liq}(1 - f_{aq}) - \delta_{gas}n_{gas}(1 - f_{aq})
\end{aligned}$$

where $x$ is cellular density (whose dimension is biomass/volume) and $n_{aq}$ and $n_{gas}$ are the number of moles in the aqueous and gas phases respectively. The parameters $g_{max}$, $k_{cap}$, $r$, $c$, and $\delta$ have the same meanings as in section 1. The new parameters are specified below :

- $k_{aq}$ is the "Monod constant", i.e. the concentration of aqueous $H_2S$ at which growth is half-maximal.
- $V_{gas}$ and $V_{liq}$ are the volumes of the gas and liquid components of the culture tube.
- $f_{aq}$ is the fraction of $H_2S$ in the aqueous phase and $(1 - f_{aq})$ is the fraction in the gas phase.
- $\delta_{gas}$ is the rate at which gaseous $H_2S$ is lost from the culture tube.

To eliminate the need to explicitly model both the aqueous and gas phases, we define the *effective* $H_2S$ *concentration*, $s_{eff} = (n_{aq} + n_{gas})/V_{liq}$, which is the concentration of $H_2S$ that one would observe if all of the $H_2S$ were in the aqueous phase. Substituting this into our equation for dynamics of cellular density, we have:

$$\begin{aligned}
\frac{dx}{dt} &= \frac{g_{max}s_{eff}f_{aq}}{k_{aq} + s_{eff}f_{aq}}\left(1 - \frac{x}{k_{cap}}\right)x \\
&= \frac{g_{max}s_{eff}}{k_{eff} + s_{eff}}\left(1 - \frac{x}{k_{cap}}\right)x
\end{aligned}$$

where in the second line we have made the substitution $k_{eff} = k_{aq}/f_{aq}$. Thus, $k_{eff}$ is the effective Monod constant, i.e. the value of $s_{eff}$ at which growth is half-maximal. The dynamics of the effective concentration $s_{eff}$ may be obtained as:

$$\begin{aligned}
\frac{ds_{eff}}{dt} &= \frac{dn_{aq}/dt}{V_{liq}} + \frac{dn_{gas}/dt}{V_{liq}} \\
&= rx - \frac{dx}{dt}c - \delta_{gas}(1 - f_{aq})s_{eff} \\
&= rx - \frac{dx}{dt}c - \delta_{eff}s_{eff}
\end{aligned}$$

where in the last line we have made the substitution $\delta_{eff} = \delta_{gas}(1 - f_{aq})$. We may regard $\delta_{eff}$ as the effective $H_2S$ leakage rate. For simplicity, we drop the subscript "$eff$" from $s_{eff}$, $\delta_{eff}$, and $k_{eff}$. Dropping these subscripts, we arrive at the equations quoted in section 1:

$$\begin{aligned}
\frac{dx}{dt} &= \frac{g_{max}s}{k + s}\left(1 - \frac{x}{k_{cap}}\right)x \\
\frac{ds}{dt} &= rx - \frac{dx}{dt}c - \delta s
\end{aligned}$$
